# Graph Ricci Curvatures Reveal Atypical Functional Connectivity in Autism Spectrum Disorder

**DOI:** 10.1101/2021.11.28.470231

**Authors:** Pavithra Elumalai, Yasharth Yadav, Nitin Williams, Emil Saucan, Jürgen Jost, Areejit Samal

**Author notes:** **Correspondence:** Nitin Williams, Areejit Samal.

## Abstract

While standard graph-theoretic measures have been widely used to characterize atypical resting-state functional connectivity in autism spectrum disorder (ASD), geometry-inspired network measures have not been applied. In this study, we apply Forman-Ricci and Ollivier-Ricci curvatures to compare networks of ASD and typically developing individuals (N = 1112) from the Autism Brain Imaging Data Exchange I (ABIDE-I) dataset. We find brain-wide and region-specific ASD-related differences for both Forman-Ricci and Ollivier-Ricci curvatures, with region-specific differences concentrated in Default Mode, Somatomotor and Ventral Attention networks for Forman-Ricci curvature. We use meta-analysis decoding to demonstrate that brain regions with curvature differences are associated to those cognitive domains known to be impaired in ASD. Further, we show that brain regions with curvature differences overlap with those brain regions whose non-invasive stimulation improves ASD-related symptoms. These results suggest the utility of graph Ricci curvatures in characterizing atypical connectivity of clinically relevant regions in ASD and other neurodevelopmental disorders.

## INTRODUCTION

Autism spectrum disorder (ASD) is an umbrella term for a diverse group of clinical presentations of neurodevelopmental disorders such as Autism, Asperger’s syndrome, childhood disintegrative disorder and pervasive developmental disorder not otherwise specified (PDD-NOS) (“National Institute of Neurological Disorders and Stroke. Autism Spectrum Disorder Fact Sheet,” 2020). ASD is characterized by difficulties in social interaction, speech and non-verbal communication, restrictive/repetitive behaviors and varying levels of intellectual disability, and can also be accompanied by neurological or psychiatric disorders (Lord et al., 2020; “National Institute of Neurological Disorders and Stroke. Autism Spectrum Disorder Fact Sheet,” 2020). Memory and movement impairments are also identified in ASD (Kristen et al., 2014; Habib et al., 2019; Zampella et al., 2021). Being highly heritable (Lord et al., 2020; Wang et al., 2017), the prevalence of ASD is globally increasing, affecting 1 in 54 children aged 8 years in the United States (Maenner et al., 2020) and 1 in 100 children aged under 6 years in India (Arora et al., 2018). In 1990, ASD was declared as a disability in the United States. While an early diagnosis is key for early intervention, an accurate and effective diagnosis of ASD is crucial (Fein et al., 2013). In order to provide a proper diagnosis and to better characterize the disorder, several studies have been undertaken to understand the pathophysiology and neurobiology of ASD (see *e.g.* Lord *et al*. (Lord et al., 2020) for a comprehensive review).

Neuroimaging methods like diffusion tensor imaging, magnetic resonance imaging (MRI) and functional magnetic resonance imaging (fMRI) are well recognized and enable us to understand structural and functional brain development in people with ASD compared to typical development and to identify the disrupted neural mechanisms underlying ASD (Clements et al., 2018; Langen et al., 2014; Solso et al., 2016; Woodward and Cascio, 2015). It also provides a means to validate clinical symptoms and cognitive theories of ASD neurobiologically (Hull et al., 2017; Langen et al., 2014; Lord et al., 2020). fMRI captures activations in different regions of the brain through the changes in blood oxygen levels (BOLD signals), and the temporal correlations between these BOLD signals are referred to as functional connectivity in the brain (Logothetis, 2008). Distant regions in the brain are activated synchronously even during rest (Biswal et al., 1995; Raichle et al., 2001) and they form the resting-state functional connectivity of the brain. Resting-state functional MRI (rs-fMRI) studies that require participants to look at a blank screen with no task demands have been used to study resting-state functional connectivity in the human brain, and have been demonstrated to be a convenient paradigm to identify neuronal correlates of neuropsychiatric disorders such as ASD (Hull et al., 2017; Woodward and Cascio, 2015). Alongside individual studies, data sharing initiatives like the Autism Brain Imaging Data Exchange (ABIDE) have offered large datasets of rs-fMRI images, encouraging and accelerating research on ASD (Di Martino et al., 2014; Hull et al., 2017; Lord et al., 2020).

Graph theory and network analysis provide objective, data-driven measures to analyze the topological architecture and connectivity patterns (human ‘connectome’) in the human brain (Bullmore and Sporns, 2009; Farahani et al., 2019; Hull et al., 2017; Rubinov and Sporns, 2010; Sporns, 2013; Van Essen et al., 2012), and can provide us with deeper insights about the functional, structural and causal organization of the brain (Farahani et al., 2019). Remarkably, many previous studies on ASD have utilized graph-theoretic analysis of rs-fMRI functional connectivity networks (FCNs) to differentiate ASD from typical development (Anderson et al., 2013; Chen et al., 2021; Di Martino et al., 2014; Harlalka et al., 2018; Itahashi et al., 2014; Keown et al., 2017; Ray et al., 2014; Redcay et al., 2013; Rudie et al., 2013; You et al., 2013), and furthermore, some of these studies (Anderson et al., 2013; Di Martino et al., 2014; Harlalka et al., 2018; Keown et al., 2017) made use of the ABIDE-I dataset. These studies investigated network characteristics such as small-worldness, modularity, clustering, efficiency, rich club organization and connection densities of the FCNs in ASD versus typical development, and reported atypical functional organization in ASD both globally and at the level of individual brain regions.

In recent years there has been an increasing interest in the development of geometric tools for analyzing complex networks (Boguñá et al., 2021), which enables the study of higher-order correlations in networks beyond pairwise interactions (Bianconi, 2021; Iacopini et al., 2019; Kartun-Giles and Bianconi, 2019). A fundamental concept in geometry is Ricci curvature (Jost, 2017), which quantifies the extent to which a space differs from being flat. Various nonequivalent definitions of graph Ricci curvature have been proposed (Chow and Luo, 2003; Forman, 2003; Ollivier, 2007; Samal et al., 2018; Sreejith et al., 2016) with an aim to capture the key properties of the classical Ricci curvature. Different notions of graph Ricci curvature have found applications in diverse areas, such as differentiating gene co-expression networks of cancer cells and healthy cells (Sandhu et al., 2015), identifying crashes and bubbles in financial networks (Samal et al., 2021; Sandhu et al., 2016), and community detection in complex networks (Ni et al., 2019; Sia et al., 2019). Ollivier-Ricci curvature (ORC) (Ollivier, 2007) and Forman-Ricci curvature (FRC) (Forman, 2003; Sreejith et al., 2016) are two widely-used notions of graph Ricci curvature.

Notably, graph Ricci curvatures have also been applied to structural and functional connectivity networks of the human brain. Farooq *et al*. (Farooq et al., 2019) applied ORC to brain structural connectivity networks to identify robust and fragile brain regions in healthy subjects. They also show that ORC can be used to identify changes in brain structural connectivity related to ASD and healthy aging. Simhal *et al*. (Simhal et al., 2020) used ORC to measure changes in brain structural connectivity of individuals with ASD before and after the infusion of autologous umbilical cord blood. ORC has also been used to study differences in brain structural connectivity networks of cognitively impaired and non-impaired multiple sclerosis patients (Farooq et al., 2020). Recently, Chatterjee *et al*. (Chatterjee et al., 2021) used a version of FRC to determine the changes in brain functional connectivity related to attention deficit hyperactivity disorder (ADHD). Additionally, FRC has been used to analyze task-based fMRI data (Weber et al., 2019) as well as to predict the intelligence of healthy human subjects (Lohmann et al., 2021). Most of these studies have also contrasted graph Ricci curvatures with standard network measures such as clustering coefficient and node betweenness centrality, and showed that graph Ricci curvatures can provide new information about brain connectivity organization. However, a systematic evaluation of the ability of graph Ricci curvatures to characterize atypical brain functional connectivity in ASD and other neurodevelopmental disorders is lacking.

In the present work, we expand the scope of curvature-based analysis for characterizing brain connectivity, by systematically applying graph Ricci curvatures to study atypical functional connectivity network organization in ASD. For this purpose, we utilized raw resting-state fMRI images of 1112 subjects from the ABIDE-I dataset and obtained FCNs for each subject by implementing a uniform preprocessing pipeline and thorough quality assessment (QA) checks. We employ FRC and ORC to compare the FCNs of individuals with ASD relative to typically developing individuals (TD), and evaluate the role of these curvature measures as indicators of atypical functional connectivity in ASD. We analyzed the brain-wide changes in FCNs by comparing average edge curvatures across the two groups, and analyzed the region-specific changes in FCNs by comparing node curvatures across the two groups. We then performed two analyses to assess the agreement of our results with relevant prior neuroimaging literature. First, we used meta-analysis decoding with respect to a large database of fMRI studies, to determine if those regions showing curvature differences are also associated to those cognitive domains known to be impaired in ASD, *e.g.* social cognition. Second, we determined if those regions showing curvature differences overlapped with those regions whose non-invasive stimulation with transcranial magnetic stimulation (TMS) (Hallett, 2007) and transcranial direct current stimulation (tDCS) (Nitsche et al., 2008) are reported in the literature to result in improvement of ASD-related symptoms.

## RESULTS

The primary goal of this study is to evaluate the utility of two notions of graph Ricci curvature, namely Forman-Ricci curvature (FRC) and Olivier-Ricci curvature (ORC), that have been recently ported to the domain of complex networks, as indicators of atypical topological organization in resting state functional connectivity networks (FCNs) of individuals with ASD. For this purpose, we analyzed spatially and temporally preprocessed rs-fMRI images of 395 individuals with ASD and 425 TD individuals from the ABIDE-I dataset as described in **STAR methods**. The demographic and clinical information for these subjects is summarized in **Table 1**. **Figure 1** is a schematic summarizing the processing pipeline for rs-fMRI data used in this study. Furthermore, **Supplementary Table S1** gives detailed information on the quality assessment and exclusion criteria for rs-fMRI dataset. Subsequently, 200 regions of interest (ROIs) or nodes were defined in the brain using the Schaefer atlas, and a 200 × 200 functional connectivity (FC) matrix was generated for each subject by computing the Pearson correlation coefficient between the time-series of all pairs of nodes. Thereafter, by combining maximum spanning tree (MST) and sparsity-based thresholding, we constructed FCNs over a wide range of graph densities between 0.02 or 2% edges and 0.5 or 50% edges, with an increment of 0.01 or 1% edges (see **STAR methods**). In a nutshell, we generated and analyzed 49 FCNs for each of the 820 subjects in the ABIDE-I dataset considered in this study.

**Figure 1:**
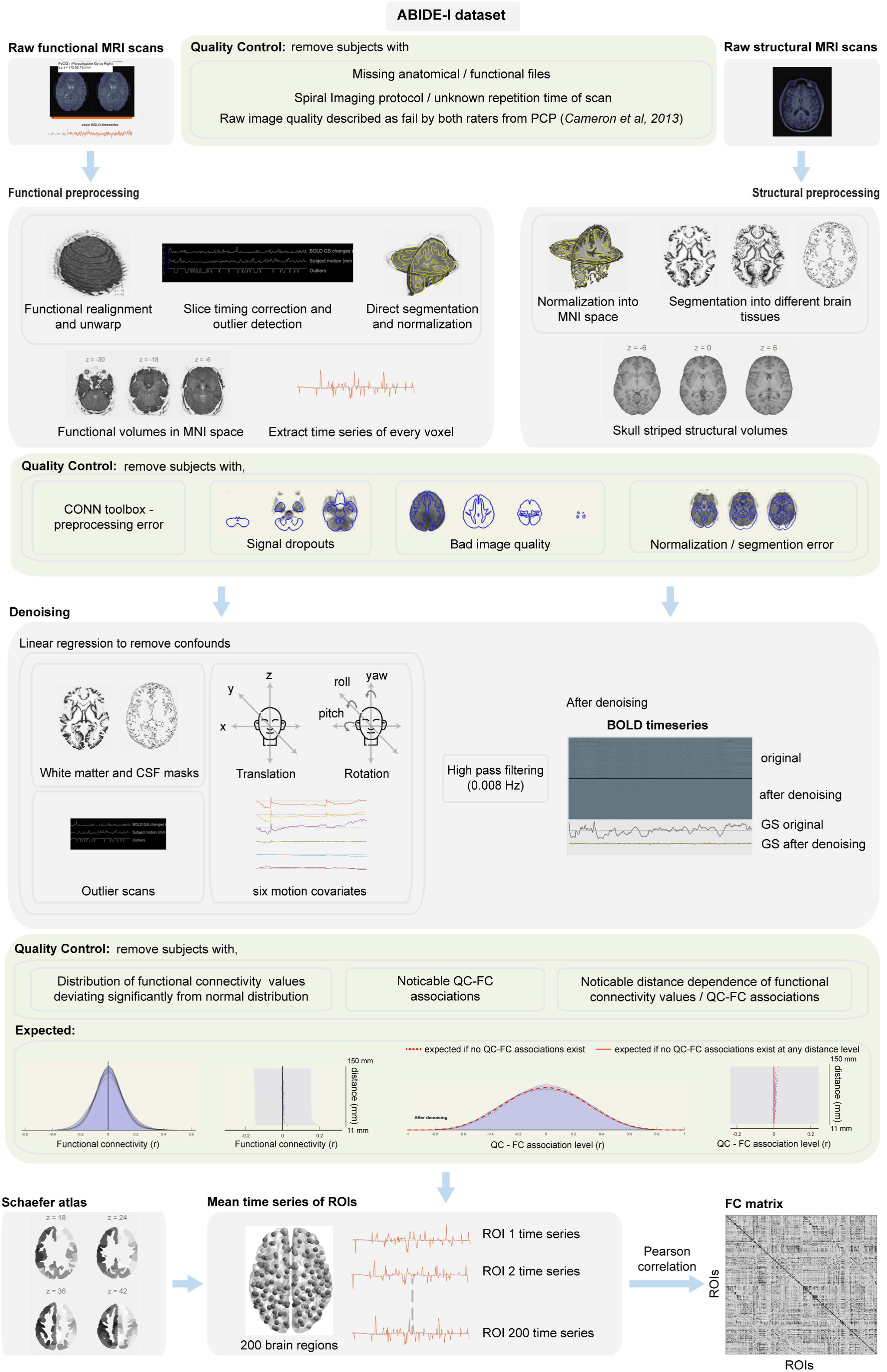
Schematic diagram summarizing the rs-fMRI processing pipeline employed in this study. The raw fMRI scans undergo four steps in spatial preprocessing, namely, motion correction, slice-timing correction, outlier detection, and direct segmentation and normalization. The raw structural MRI scans are normalized to the Montreal Neurological Institute (MNI) space, and segmented into grey matter, white matter and cerebrospinal fluid (CSF) areas. In the temporal preprocessing or denoising step, the BOLD time series of each voxel is extracted and the remaining physiological and motion confounds are removed using linear regression. The confounds include white matter and CSF masks, subject-motion parameters and outlier scans. The residual BOLD time series of each voxel undergoes a high-pass filtering at 0.008 Hz. The Schaefer atlas is used to parcellate the brain into 200 regions of interest (ROIs) and the mean time series for each ROI is computed. Finally, Pearson correlation coefficient is computed between all pairs of ROIs, resulting in a 200 × 200 functional connectivity (FC) matrix. Thorough quality assessment (QA) checks were implemented both before and after preprocessing. In this figure, the head icon under denoising section is made by Freepik from flaticon.com (https://www.flaticon.com/authors/freepik).

**Table 1.** Summary of demographic and clinical information for the 820 subjects from ABIDE-I project that fulfil the inclusion criteria and were selected for network analysis in this study. 395 subjects belong to the autism spectrum disorder (ASD) group and 425 subjects belong to the typically developing (TD) group. The subjects in both groups are age-matched (*p* = 0.835). Handedness data were missing for 137 ASD and 148 TD subjects. ADI-R social data were missing for 120 ASD participants. ADI-R verbal data were missing for 119 ASD participants.

| Characteristics | ASD Group | TD Group |
| --- | --- | --- |
| **number of subjects** | 395 (44 female) | 425 (78 female) |
| **age (in years)** |  |  |
| mean ± s.d. | 15.6 ± 7.1 | 15.51 ± 6.23 |
| range | 7 - 58 | 6 - 57 |
| **handedness (*n*)** |  |  |
| left | 29 | 27 |
| right | 225 | 247 |
| mixed | 3 | 3 |
| ambidextrous | 1 | 0 |
| **ADI-R Social** |  |  |
| mean ± s.d. | 19.72 ± 5.25 | - |
| range | 7 - 30 | - |
| **ADI-R Verbal** |  |  |
| mean ± s.d. | 15.95 ± 4.25 | - |
| range | 2 - 26 | - |

### Brain-wide changes in functional connectivity networks

To investigate the differences in the global organization of FCNs between the ASD and TD groups, we computed the average edge FRC and average edge ORC across the 49 FCNs across the graph densities 2% - 50% for each subject. To compare the average edge curvatures at each graph density between the ASD and TD groups, we employed a two-tailed two-sample t-test followed by FDR correction (see **STAR methods**). In **Figures 2a** and **2b**, we show the differences in average edge FRC and average edge ORC, respectively, between the ASD and TD groups across the graph densities 2% - 50%. We find that average edge FRC is significantly lower (*p* < 0.05, FDR-corrected) in the ASD group compared to the TD group in the graph density range 5%-50% (**Figure 2a**). Similarly, we find that average edge ORC is lower (*p* < 0.05, FDR-corrected) in the ASD group compared to the TD group albeit the differences were insignificant (*p* > 0.05, FDR-corrected) in the graph density ranges 2%-5% and 24%-33% (**Figure 2b**). Although the directionality of the differences with the two discrete Ricci curvatures is the same for the two groups, that is, average edge curvature in the ASD group is lower than that in the TD group, it is important to emphasize that the two discrete Ricci curvatures capture different aspects of the classical Ricci curvature, and thus, cannot serve as alternative measures across different types of networks. Specifically, ORC captures the volume growth property of the classical Ricci curvature whereas FRC captures the geodesic dispersal property (Samal et al., 2018). While ORC has a deeper correspondence with the classical Ricci curvature, FRC is based on a simple combinatorial expression which is significantly faster to compute in larger networks. After comparing the average edge curvatures between FCNs of ASD and TD groups, we find that the statistical test (t-test followed by FDR correction) yielded lower p-values after FDR correction for average edge FRC compared to average edge ORC across most of the considered graph densities (**Supplementary Table S2**). In other words, the differences between FCNs for the two groups are more pronounced for the average edge FRC than average edge ORC.

**Figure 2:**
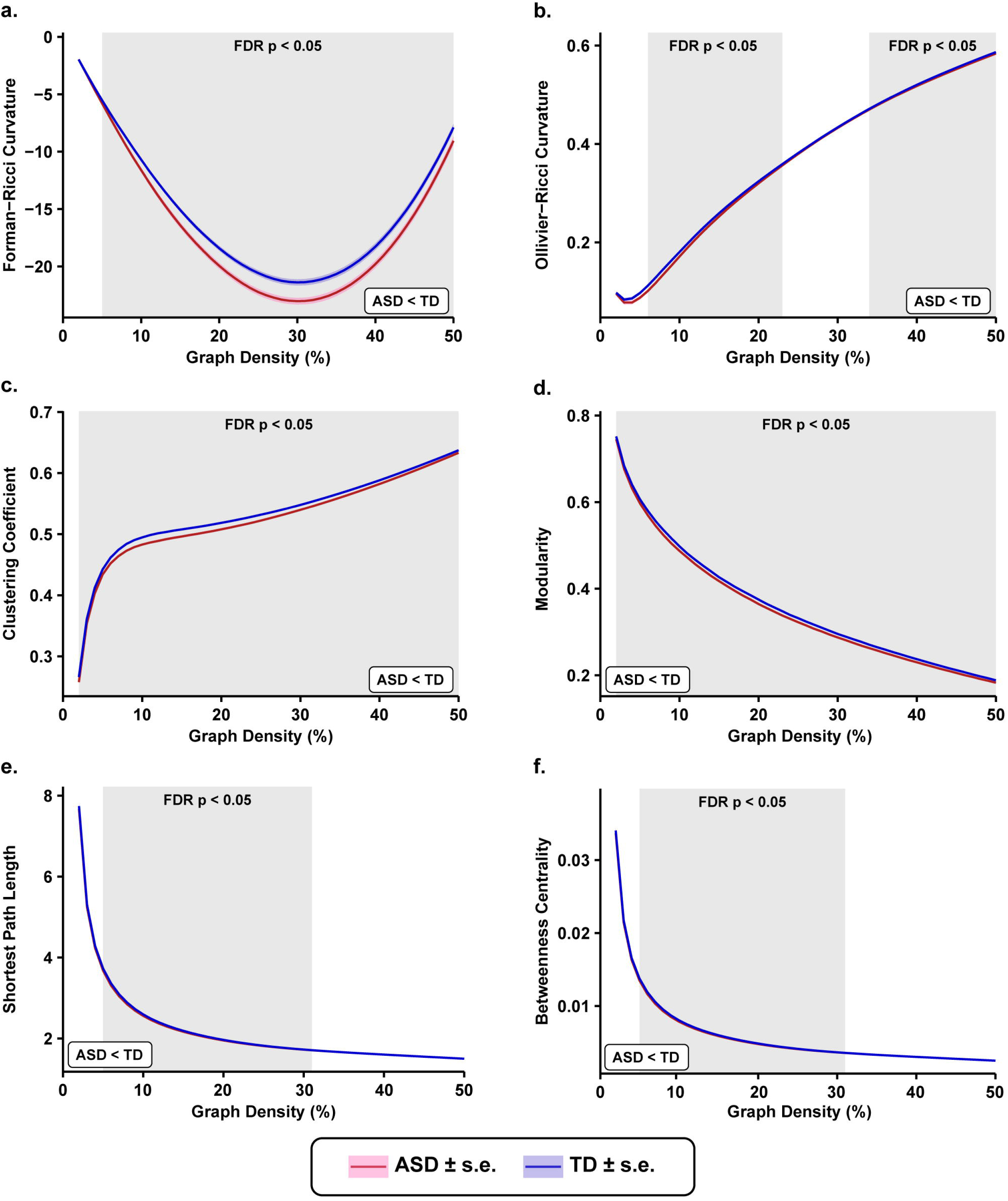
Brain-wide changes in functional connectivity networks. Comparison plots of global changes in functional connectivity networks (FCNs) as captured by network measures between 395 subjects with autism spectrum disorder (ASD) and 425 age-matched typically developing individuals (TD). Each network measure was compared over a wide range of graph densities between 0.02 (i.e., 2% edges) and 0.5 (i.e., 50% edges), with an increment of 0.01 (i.e., 1% edges). The shaded regions in each plot indicate statistically significant differences (*p* < 0.05, FDR-corrected) between the two groups at the corresponding graph densities on the x-axis. Even though the differences are not explicit from the plots (e) and (f), the directionalities are programmatically verified. (a) Average Forman-Ricci curvature (FRC) of edges is significantly reduced in the ASD group across graph densities 5% - 50%. It is evidently the most visually observable difference among all the other network measures studied. (b) Average Ollivier-Ricci curvature (ORC) of edges is significantly reduced in the ASD group across graph densities 6% - 23% and 34% - 50%. (c) Average clustering coefficient is significantly reduced in the ASD group across all graph densities (2% - 50%). (d) Modularity is significantly reduced in the ASD group across all graph densities (2% - 50%). (e) Average shortest path length is significantly reduced in the ASD group across graph densities 5% - 31%. (f) Average node betweenness centrality is significantly reduced in the ASD group across graph densities 5% - 31%.

To gain a deeper understanding of the altered global organization of FCNs between the ASD and TD groups, we also compared six other global network measures, namely, average clustering coefficient, modularity, average shortest path length, average node betweenness centrality, global efficiency and average local efficiency. We find that the average clustering coefficient is significantly lower (*p* < 0.05, FDR-corrected) in the ASD group compared to the TD group in the graph density range 2%-50% (**Figure 2c**). Moreover, our results for clustering coefficient are consistent with results from previous studies that have employed graph-theoretic measures to analyze resting state FCNs in ASD (Harlalka et al., 2018; Itahashi et al., 2014; Rudie et al., 2013). Thereafter, we find that the modularity of the FCNs is significantly reduced in the ASD group compared to the TD group in the graph density range 2%-50% (**Figure 2d**), and our results are consistent with the results from previous studies (Harlalka et al., 2018; Rudie et al., 2013). Further, we find that the average shortest path length of the FCNs is significantly lower in the ASD group compared to the TD group in the graph density range 5%-31% (**Figure 2e**), and our results are consistent with results from previous studies (Itahashi et al., 2014; Rudie et al., 2013). Lastly, we find that average node betweenness centrality is significantly lower in the ASD group compared to the TD group in the density range 5% - 31% (**Figure 2f**).

Furthermore, we have computed two global measures that characterize how efficiently information is exchanged within a network, namely global efficiency and average local efficiency. We find that global efficiency is significantly higher (*p* < 0.05, FDR-corrected) in the ASD group compared to the TD group in the graph density range 4% - 31% (**Supplementary Figure S1a**). Note that the direction of the effects observed for global efficiency is opposite to the direction of effects observed for average shortest path length (**Figure 2e**), as global efficiency is defined as the average of reciprocal shortest path lengths between all pairs of nodes in a network. Moreover, our results for global efficiency are consistent with the results from previous studies (Harlalka et al., 2018; Itahashi et al., 2014; Rudie et al., 2013). We find that average local efficiency is significantly lower in the ASD group compared to the TD group in the graph density range 2%-50% (**Supplementary Figure S1b**). Note that the results for average local efficiency are similar to the results of average clustering coefficient (**Figure 2c**), since the two network measures are closely related to each other. Moreover, our results for average local efficiency are consistent with results from previous studies (Harlalka et al., 2018; Itahashi et al., 2014; Rudie et al., 2013).

### Region-specific changes in functional connectivity networks

Given the significant differences in FRC and ORC of the entire brain between the ASD group and the TD group, we evaluated node-level curvature differences in the FCNs, and determined how these differences are distributed across the 7 resting state networks (RSNs) in the brain. For this purpose, we first computed node FRC and node ORC for all the 200 nodes, across the 49 FCNs with graph densities 2% - 50% for each subject. Second, to identify the set of nodes that show significant differences between the ASD and TD groups, we compared the area under the curve (AUC) of the node FRC and the node ORC for each node using a two-tailed two-sample t-test followed by FDR correction (see **STAR methods**).

In **Figure 3** and **Supplementary Figure S2**, we show the nodes or regions that exhibit significant differences (*p* < 0.05, FDR-corrected) in FRC and ORC, respectively, between the ASD and TD groups. We identify 83 regions that show significant between-group differences in FRC and 14 regions that show significant between-group differences in ORC. For FRC, the significant regions are spread across the 7 RSNs. However, they are mainly concentrated within 3 RSNs namely, default network (26 significant regions), somatomotor network (30 significant regions) and salient ventral attention network (13 significant regions). In the default network, RH_Default_pCunPCC_2 (7, -49, 31), RH_Default_PFCdPFCm_6 (28, 30, 43) and LH_Default_pCunPCC_2 (−5, -55, 27) showed the lowest FDR corrected p-values. In the somatomotor network, LH_SomMot_7 (−47, -9, 46), RH_SomMot_7 (58, -5, 30) and LH_SomMot_10 (−39, -24, 58) showed the lowest FDR corrected p-values. In the salient ventral attention network, RH_SalVentAttn_TempOccPar_2 (60, -38, 17), RH_SalVentAttn_Med_3 (9, 4, 65) and RH_SalVentAttn_PrC_1 (51, 4, 40) showed the lowest FDR corrected p-values. For ORC, the significant regions are concentrated within the 2 RSNs namely, default network and somatomotor network. In the default network, regions LH_Default_Temp_3 (−56, -6, -12), LH_Default_PFC_4 (−13, 63, -6), and RH_Default_Temp_5 (52, -31, 2) exhibited the lowest FDR corrected p-values. In the somatomotor network, the region LH_SomMot_3 (−37, -21, 15) exhibits the lowest FDR corrected p-value. Detailed information about the abbreviations of region names can be found at: https://github.com/ThomasYeoLab/CBIG/tree/master/stable_projects/brain_parcellation/Schaefer2018_LocalGlobal (Schaefer et al., 2018). Thus, our node-level results for FRC and ORC suggest that the nodes or brain regions showing significant differences were not distributed evenly across the 7 RSNs, but concentrated within the default network, somatomotor network and salient ventral attention network.

**Figure 3:**
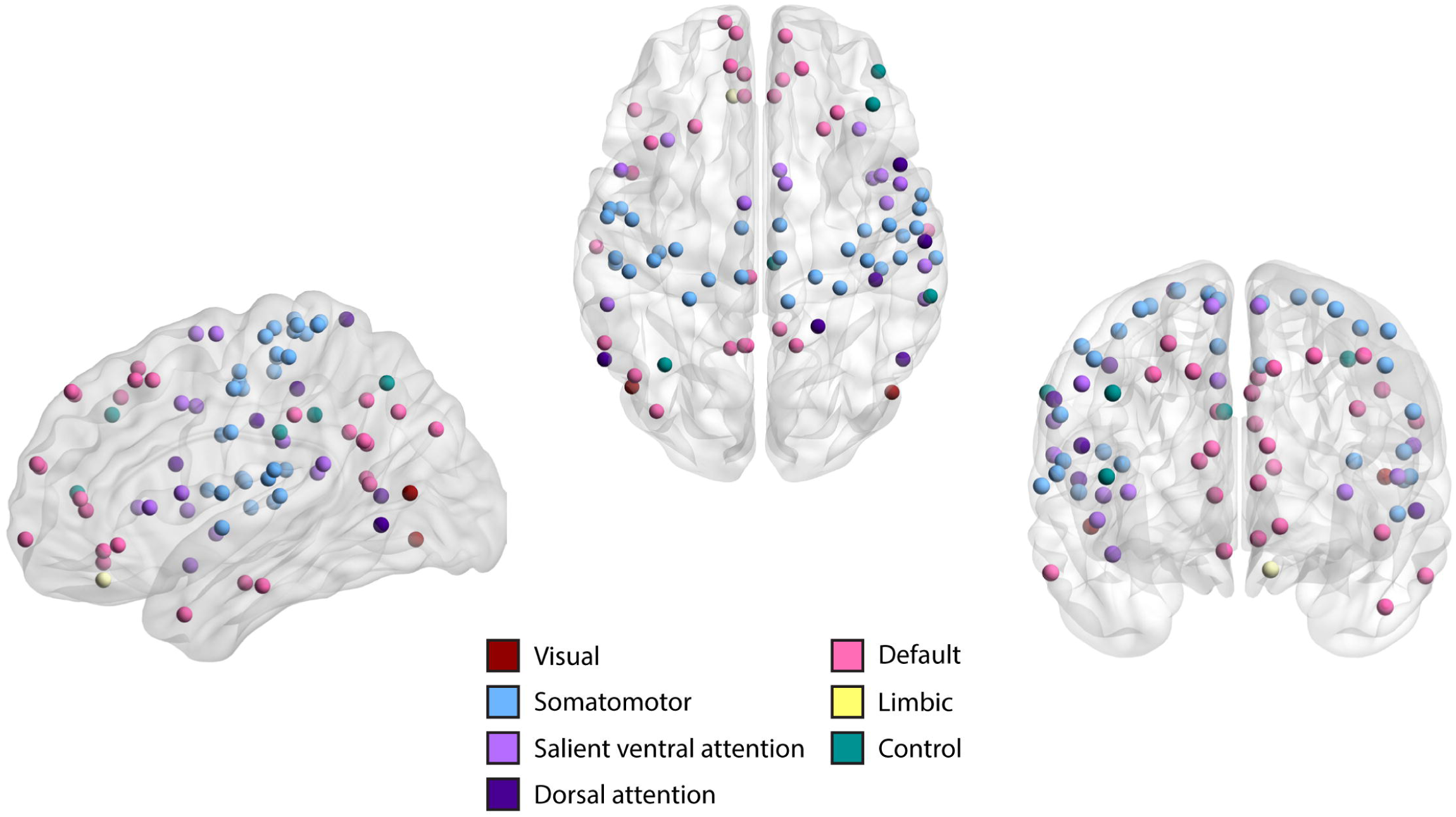
Region-specific changes in functional connectivity networks. Visual representation of 83 nodes or regions in the brain that are significantly different (*p* < 0.05, FDR-corrected) between individuals with autism spectrum disorder (ASD) and typically developing individuals (TD), as captured by Forman-Ricci curvature (FRC) of the nodes in the functional connectivity networks (FCNs) of the subjects. The nodes are defined using the Schaefer atlas and each node belongs to one of 7 resting state networks (RSNs) as listed in the figure legend. We find that identified nodes are mainly concentrated within the default network, somatomotor network, and salient ventral attention network. This figure was created using BrainNet Viewer (Xia et al., 2013).

After applying both FRC and ORC to resting state FCNs in ASD and TD groups, we find that the between-group differences in FRC of the FCNs are more pronounced compared to ORC, both at the global-level and the node-level. Therefore, we mainly focus on the nodes identified using FRC in further analyses. We reiterate that the two discrete Ricci curvatures capture different aspects of the classical Ricci curvature, and thus, neither of the two measures can be treated as an alternative to the other.

### Agreement of results from node-level network analysis to fMRI literature

We assessed the agreement of our results on ASD-related region-specific differences in FRC to relevant previous neuroimaging literature. One form of validation we performed was to determine if those regions showing significant ASD-related differences in FRC were also associated in the fMRI literature, to cognitive domains impaired in ASD, *e.g.* social cognition. To do this, we first partitioned the set of significant brain regions according to their respective RSNs and determined the cognitive domains associated to the significant regions in each RSN using Neurosynth meta-analysis (see **STAR methods**). The Neurosynth analysis enables identifying the cognitive domains associated to the significant regions in an RSN more rigorously than just assuming it as the putative functional role of that RSN. The first step in the Neurosynth analysis involves identifying terms relating to cognition, perception and behavior for each significant brain region in a given RSN. The second step involves calculating the frequency counts for all the terms in an RSN. The third step involves thresholding the frequency counts of these terms with respect to the frequency counts associated with equivalent null models (see **STAR methods**).

We limited the Neurosynth analysis only to default network, somatomotor network and salient ventral attention network, since a considerable number of regions are detected in these RSNs. The high number of regions with significantly different FRC in these RSNs makes the interpretation of results from these RSNs more robust to the occurrence of false positives. Another reason for considering the above-mentioned RSNs is that the regions identified in these RSNs are nearly bilaterally symmetrical. **Figure 4** shows the significant brain regions separately for each of the three RSNs and the associated word clouds highlighting the behavioral relevance of the significant regions in each RSN. **Supplementary Table S3** lists the significant brain regions and the terms associated with all seven RSNs.

**Figure 4:**
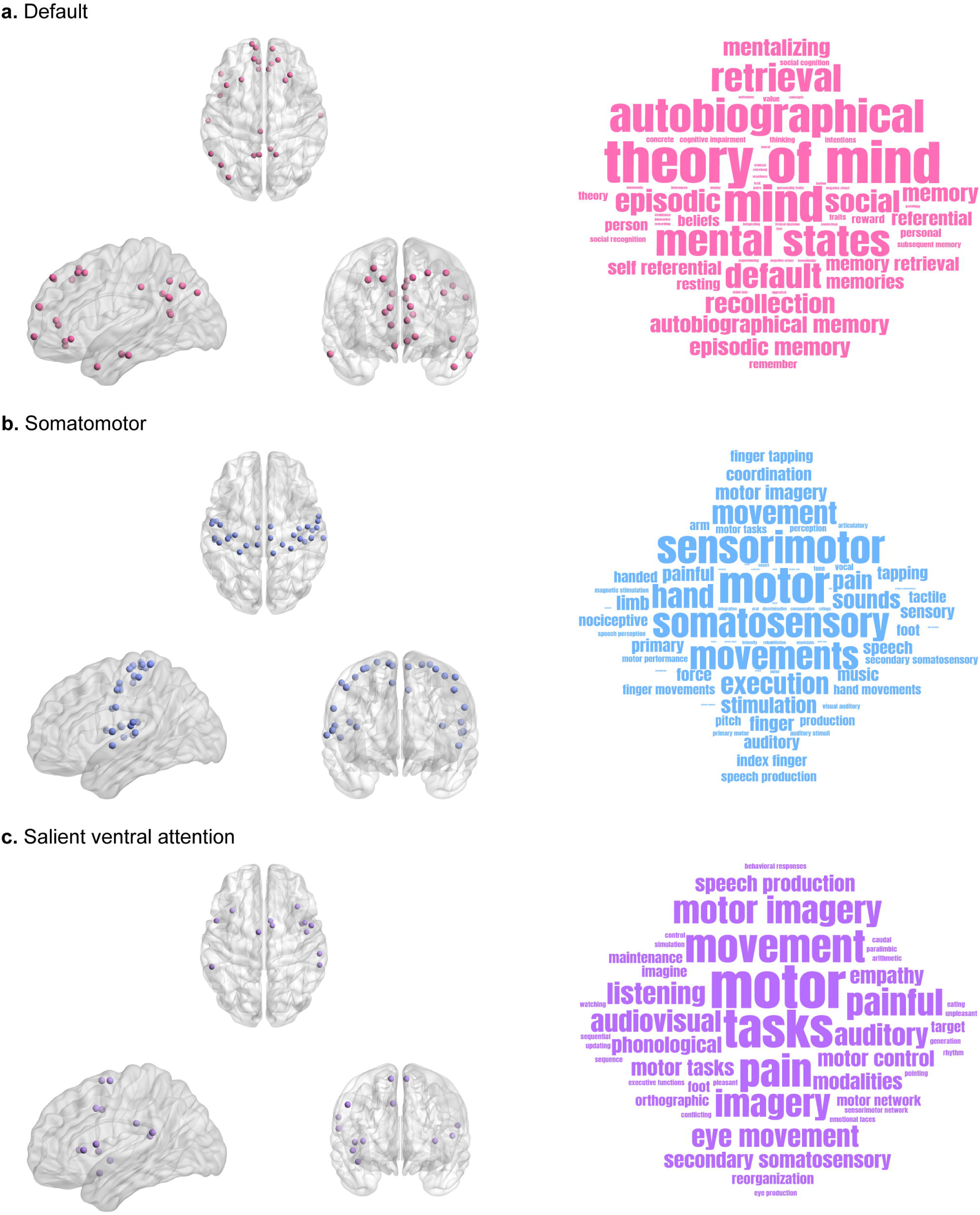
Agreement of results from node-level network analysis to fMRI literature. Visual representations of nodes or regions in different resting state networks (RSNs) that are significantly different (*p* < 0.05, FDR-corrected) between individuals with autism spectrum disorder (ASD) and typically developing individuals (TD) as captured by Forman-Ricci curvature (FRC) of the nodes, and the corresponding word clouds depicting the behavioral relevance of the nodes identified in each RSN. The size of the terms in each word cloud indicates their frequency count. Note that size of the terms in each word cloud are scaled separately and thus the frequency counts cannot be compared across word clouds. (a) Nodes in the default network that show significant differences in FRC, and the corresponding word cloud. The nodes identified in the default network are associated with tasks related to social cognition and memory. (b) Nodes in the somatomotor network that show significant differences in FRC, and the corresponding word cloud. The nodes identified in the somatomotor network are associated with tasks related to movement. (c) Nodes in the salient ventral attention network that show significant differences in FRC, and the corresponding word cloud. The nodes identified in the salient ventral attention network are associated with tasks related to movement and language. In this figure, the visualizations of brain regions are created using BrainNet Viewer (Xia et al., 2013) and the word clouds are generated using wordclouds.com (https://www.wordclouds.com).

The word cloud for the default network shows terms associated with social cognition (**Figure 4a**), such as ‘theory-of-mind’, ‘social’, ‘social recognition’, ‘social cognition’, ‘person’, ‘personal ’and ‘personality traits’. Impairments in social cognition are known to be characteristic of individuals with ASD (Kasari and Patterson, 2012; Kristen et al., 2014; Senju, 2012). In the default network, we can also find terms associated with memory (**Figure 4a**) such as ‘memory’, ‘memories’, ‘memory retrieval’, ‘retrieval‘ ’recollection’, ‘remember’, ‘autobiographical memory’, ‘episodic memory ’and ‘subsequent memory’. Just as with social cognition, memory impairments are often a feature of ASD (Griffin et al., 2021; Habib et al., 2019; Kristen et al., 2014; Solomon et al., 2016), though memory impairments are not part of the standard diagnostic criteria for ASD. For the somatomotor network, we find terms associated with movement (**Figure 4b**), such as ‘motor’, ‘motor tasks’, ‘motor performance’, ‘movements’, ‘coordination’, ‘limb’, ‘arm’, ‘hand movements’, ‘handed’, ‘finger’, ‘finger movements’, ‘finger tapping’, ‘tapping’, ‘index finger’, and ‘force’. For the salient ventral attention network also, we find terms associated with movement (**Figure 4c**), such as ‘motor’, ‘motor function’, ‘motor control’, ‘eye movement’, ‘tapping’, ‘mental imagery’, ‘imagery ’and ‘mirror’. Notably, the literature reports movement impairments to characterize individuals with ASD (Bhat, 2021; Grace et al., 2018; Ming et al., 2007; Zampella et al., 2021). For the salient ventral attention network, we also find terms associated with language (**Figure 4c**), such as ‘phonological’, ‘speech’, ‘production’, ‘speech production’, ‘orthographic’, ‘articulatory’, ‘pseudo-words ’and ‘listened’. In line with this, individuals with ASD often present with speech and communication difficulties (Davidson and Ellis Weismer, 2017; Ellis Weismer et al., 2010; Pickles et al., 2009). Hence, we find that those regions exhibiting ASD-related curvature differences are also associated to those cognitive domains known to be impaired in ASD. These results serve as a form of validation, based on the fMRI literature, that FRC identifies clinically relevant brain regions underlying ASD.

We also evaluated the node-level differences in two standard network measures namely, clustering coefficient and node betweenness centrality. Notably, ORC is related to clustering in networks (Samal et al., 2018). We identify 78 brain regions that show significant differences (*p* < 0.05, FDR-corrected) in clustering coefficient, and 4 brain regions that show significant differences (*p* < 0.05, FDR-corrected) in node betweenness centrality (**Supplementary Table S4**). The brain regions identified by clustering coefficient are concentrated in three RSNs namely, default network, somatomotor network and salient ventral attention network (**Supplementary Figure S3**). Further, we computed the overlap between sets of significant brain regions identified by each of the four node-level network measures used in our study. First, we found 8 brain regions that are commonly identified by both FRC and ORC, namely, LH_SomMot_1 (−51, -5, -2), LH_SomMot_3 (−37, -21, 15), LH_Default_PFC_4 (−13, 63, -6), RH_SomMot_3 (38, -13, 14), RH_DorsAttn_Post_2 (52, -60, 9), RH_SalVentAttn_FrOperIns_2 (46, -3, -4), RH_SalVentAttn_TempOccPar_2 (60, -38, 17) and RH_Default_Temp_2 (61, -13, -21). Second, we found 71 brain regions that are commonly identified by FRC and clustering coefficient. Third, we found 5 brain regions that are commonly identified by ORC and clustering coefficient. Fourth, we found that 1 brain region is commonly identified by FRC and node betweenness centrality. Fifth, we found 2 brain regions that are commonly identified by ORC and node betweenness centrality. We do not find any brain regions that are commonly identified by clustering coefficient and node betweenness centrality.

Subsequently, we determined if there is a relationship between the FRC of brain regions that showed significant differences and clinical scores for symptom severity in ASD. To do this, we related the FRC of just the brain regions which showed significant differences in each RSN with the behavioral function associated with that RSN, as determined by the Neurosynth meta-analysis decoding. We performed this analysis with the FRC values and symptom severity only for individuals with ASD, not the TD individuals. First, we used ADI-R social score as a measure of social cognition and related this score with the FRC of regions in the default network. Second, we used ADI-R verbal score as a measure of language and related this score with the FRC of regions in the salient ventral attention network. To test the relationship between FRC values and clinical scores, we computed partial correlations with age and gender as covariates, followed by FDR correction to control the occurrence of false positives (see **STAR methods**). We chose the ADI-R scores among all the possible clinical scores because they are available for the most number of participants (*n* = 275) in the ASD group and the ADI-R social and ADI-R verbal scores are appropriate means to capture symptom severity in autism compared to other clinical scores (Lefort-Besnard *et al*., 2020). Note that we also found memory-related terms in default network (**Figure 4a**), and movement-related terms in somatomotor network (**Figure 4b**) and salient ventral attention network (**Figure 4c**). However, there was no suitable memory-related score in the ABIDE-I dataset, as memory impairments are not part of the standard diagnostic criteria for ASD. We did not include any movement-related scores in our analysis since such scores were only available for a few participants in the ASD group.

We did not find any nodes that showed significant correlations between FRC and clinical scores after FDR correction. Prior to FDR correction, FRC for the node LH_Default_Temp_1 (−47, 8, - 32) in the default network was positively correlated with ADI-R social score (*r* = 0.122, *p* = 0.044), FRC for the node LH_SalVentAttn_ParOper_1 (−56, -40, 20) in the salient ventral attention network was positively correlated with ADI-R verbal score.

We also repeated the analysis for the brain regions with significantly different clustering coefficient values in the 2 RSNs namely, default network and salient ventral attention network. Similar to FRC, these brain regions show behavioral relevance and are associated to social cognition and memory in default network, and movement and language in salient ventral attention network (**Supplementary Figure S4**). Thus, we correlated the clustering coefficient values of significant brain regions in default network with the ADI-R social score and the clustering coefficient values of significant brain regions in salient ventral attention network with the ADI-R verbal score. In this analysis for clustering coefficient, we did not find any significant correlations with both ADI-R social scores and ADI-R verbal scores after FDR correction. To sum up, neither FRC nor clustering coefficient show evidence for a relationship with symptom severity in individuals with ASD.

### Agreement of results from node-level network analysis to TMS/tDCS literature

In addition to the meta-analysis decoding, we performed one more analysis to determine the agreement of our results with relevant previous neuroimaging literature. Specifically, we determined the overlap between those brain regions showing FRC differences and those whose non-invasive stimulation using TMS or tDCS, resulted in improvement of ASD-related symptoms. To do this, we performed a literature search on PubMed to identify the set of brain regions whose non-invasive stimulation using TMS or tDCS yielded positive effects on ASD symptoms. The exact details of the PubMed search query are provided in **Table 2**. Then, we compared this set of brain regions to those with altered FRC values in resting-state fMRI FCNs of individuals with ASD. **Figure 5** summarizes the workflow we employed to collect and classify the eligible articles from the literature survey.

**Figure 5:**
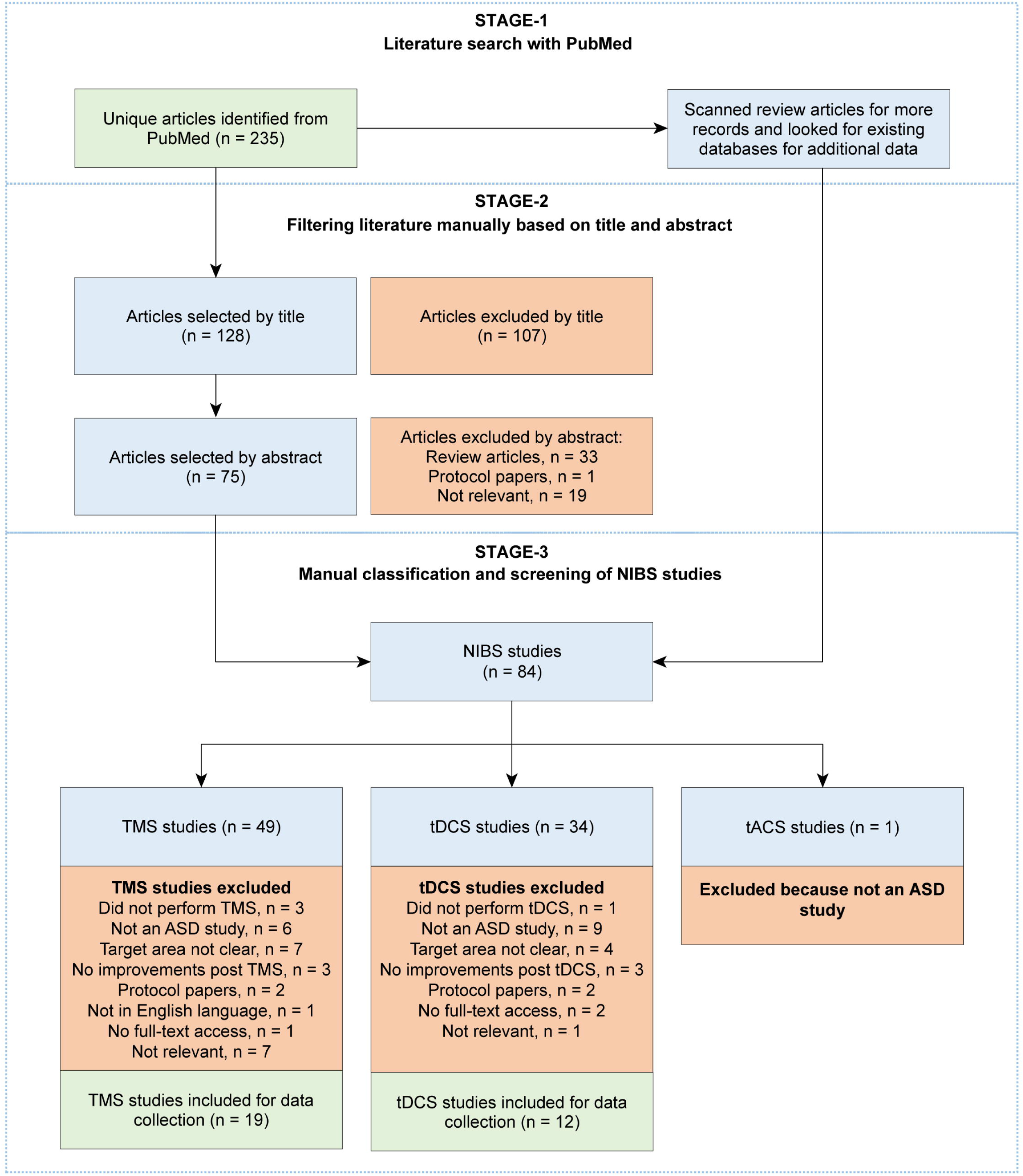
Summary of the workflow employed to compile data from non-invasive brain stimulation (NIBS) experiments. The workflow is presented according to PRISMA statement (Moher et al., 2009). First, we identified 235 potential records from PubMed. Second, we filtered the articles based on title and abstract. Third, we scanned review articles for more records and looked for existing databases for additional data. After performing the above steps, we were left with 84 potential NIBS studies. Finally, we classified the studies based on the stimulation technique (TMS/tDCS/tACS) and screened the studies individually for eligibility. We were left with 19 TMS studies and 12 tDCS studies, which were used to extract experimental data.

**Table 2.** Literature search for non-invasive brain stimulation studies in ASD. Detailed summary of the electronic search query on PubMed that we used to obtain our original corpus of articles that perform non-invasive brain stimulation on individuals with ASD.

| Search date | Search query | Search filters | Source |
| --- | --- | --- | --- |
| October 18, 2021 | ((transcranial magnetic stimulation) AND (autism)) OR ((transcranial magnetic stimulation) AND (Asperger)) OR ((transcranial magnetic stimulation) AND (PDD NOS)) OR ((transcranial direct current stimulation) AND (autism)) OR ((transcranial direct current stimulation) AND (Asperger)) OR ((transcranial direct current stimulation) AND (PDD NOS)) OR ((transcranial alternating current stimulation) AND (autism)) OR ((transcranial alternating current stimulation) AND (Asperger)) OR ((transcranial alternating current stimulation) AND (PDD NOS)) OR ((TMS) AND (autism)) OR ((TMS) AND (Asperger)) OR ((TDCS) AND (autism)) OR ((TDCS) AND (Asperger)) OR ((TACS) AND (autism)) OR ((TACS) AND (Asperger)) | year : no filter,<br>article attribute : no filter,<br>language : no filter,<br>age : no filter,<br>sex : no filter,<br>publication date: no filter | PubMed<br>(n = 235) |

The studies employing TMS have reported positive effects in ASD-related symptoms after stimulating 4 target regions, namely, premotor cortex, dorsolateral prefrontal cortex (DLPFC), pars triangularis, and pars opercularis. The studies employing tDCS have reported positive effects in ASD symptoms after stimulating 2 target regions, namely, DLPFC and left primary motor cortex. **Supplementary Tables S5** and **S6** provide a detailed summary of the TMS and tDCS studies, respectively.

Note that the target regions in these experiments are cortical regions that are defined differently from the ROIs (or nodes) defined in our study that are a part of the Schaefer 200 parcels atlas (Schaefer et al., 2018). Therefore, in order to compare the results of our node-level analysis with the effects of stimulating the target regions, we mapped the Brodmann areas that correspond to target regions (Cieslik et al., 2013; Strotzer, 2009) to the 200 Schaefer ROIs (see **STAR methods** and **Supplementary Table S7**).

Based on the data collected from previous NIBS experiments (**Supplementary Tables S5** and **S6**), we identified five target regions that show evidence for improvement in behavioral or cognitive symptoms associated with ASD following TMS or tDCS, namely, premotor cortex, pars triangularis, pars opercularis, DLPFC and left primary motor cortex. These five target regions correspond to Brodmann areas 6, 45, 44, 9, 46 and 4, respectively. Note that DLPFC comprises two Broadman areas, 9 and 46 (Cieslik et al., 2013). We found these Brodmann areas to encompass 31 ROIs (or nodes) in the Schaefer 200 parcels atlas. Out of these 31 ROIs, 18 ROIs also show significant ASD-related differences in FRC and 13 ROIs show significant ASD-related differences in clustering coefficient. None of these 31 ROIs show significant ASD-related differences in ORC or node betweenness centrality. A visual representation of these ROIs is provided in **Figure 6**. Notably, the 18 ROIs with significant ASD-related differences in FRC are a superset of the 13 ROIs with significant between-group differences in clustering coefficient. These results serve as a form of validation, based on the literature on outcomes from TMS and tDCS experiments, that FRC identifies clinically relevant brain regions underlying ASD. Further, FRC identifies some regions that might be clinically relevant in ASD, but are not identified by other node-based network measures, *e.g.* clustering coefficient. **Table 3** lists the target regions that show improvement in clinical symptoms associated with ASD following TMS or tDCS, the corresponding ROIs in the Schaefer atlas that show significant between-group differences in node-level network measures, the network measure that captured the differences, and the experimental studies that report the effects.

**Figure 6:**
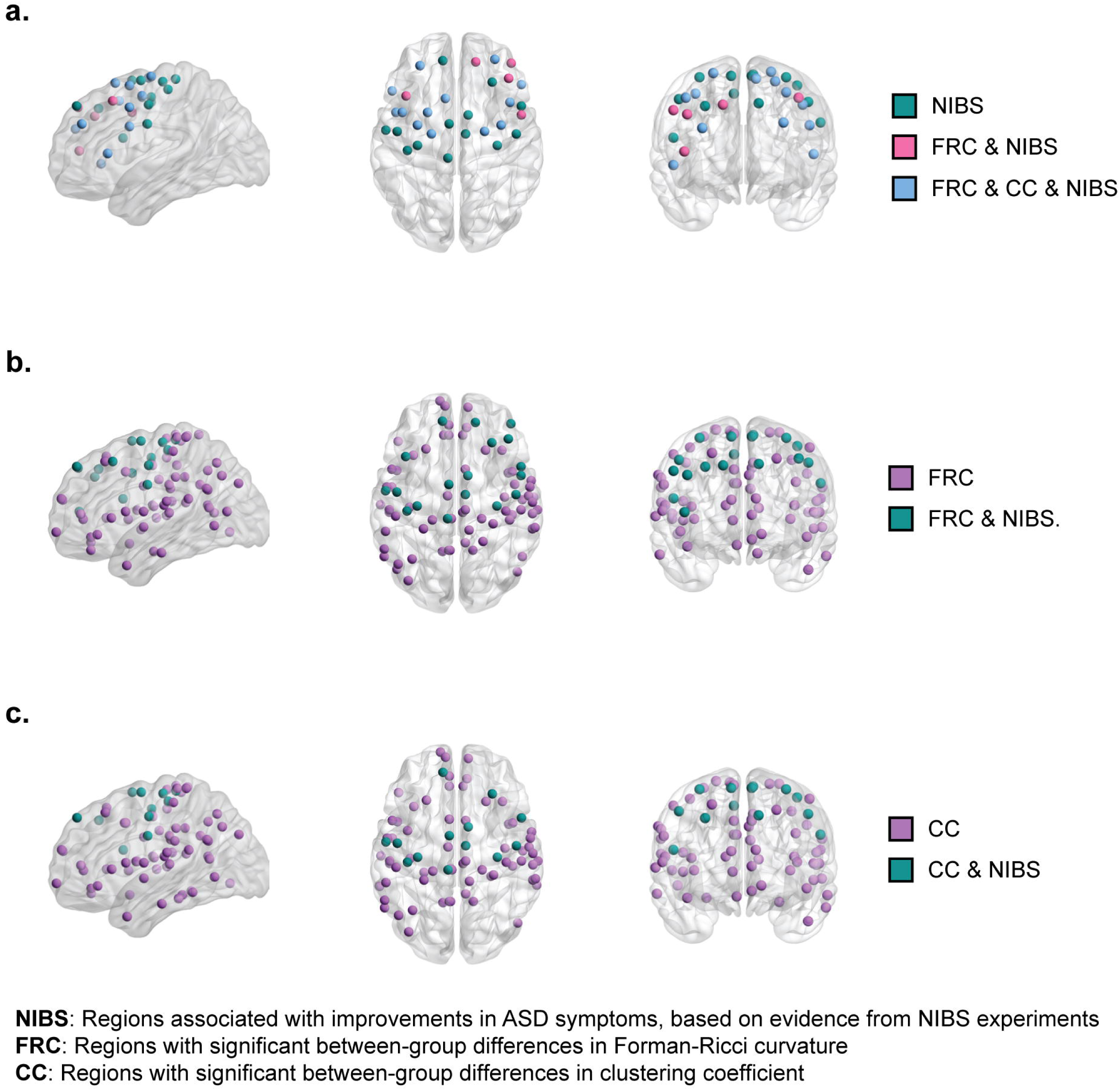
Agreement of results from node-level network analysis to TMS/tDCS literature. Visual representation of nodes or regions with significant between-group differences in node-level network measures that exhibit improvements in clinical symptoms of ASD when stimulated, based on evidence from published NIBS experiments on subjects with ASD. (a) We found 31 nodes with experimental evidence out of which 13 nodes are identified by both FRC and clustering coefficient and 5 nodes are identified only by FRC. (b) 83 nodes identified by FRC out of which 18 nodes have experimental evidence. (c) 78 nodes identified by clustering coefficient out of which 13 nodes have experimental evidence. The visualizations of the brain regions are created using BrainNet Viewer (Xia et al., 2013).

**Table 3.**
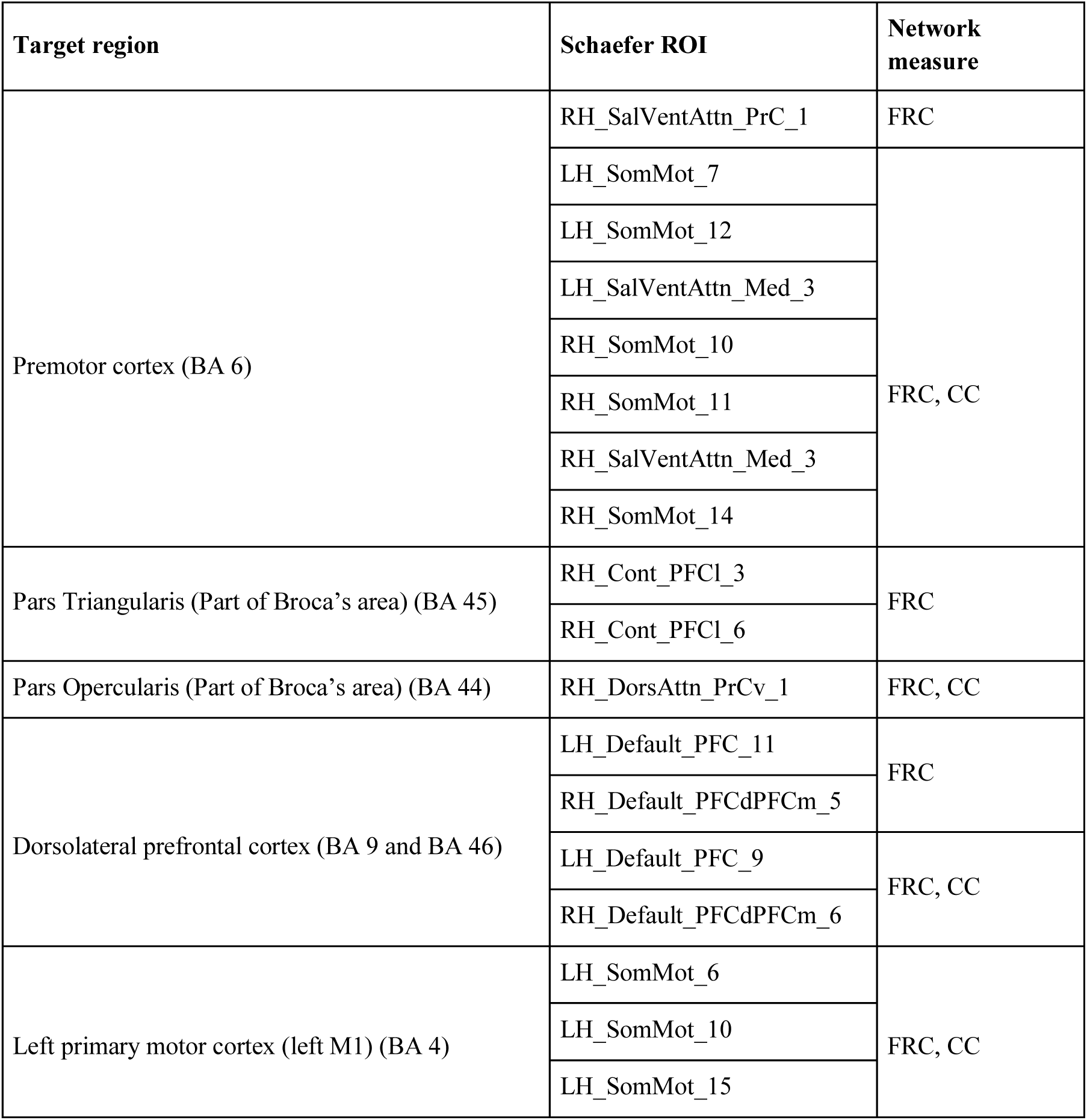
Agreement of results from node-level network analysis to TMS/tDCS literature. The list of the target brain regions that show improvement in clinical symptoms associated with ASD following TMS or tDCS procedure, the corresponding ROIs in the Schaefer atlas that show significant between-group differences in node-level network measures and the network measures that capture the differences (Forman-Ricci curvature (FRC), clustering coefficient (CC)).

## DISCUSSION

Graph Ricci curvatures have not been previously applied to study atypical resting-state functional connectivity in ASD. In the present work, we used two notions of graph Ricci curvature, namely Forman-Ricci curvature (FRC) and Ollivier-Ricci curvature (ORC) to compare the resting-state FCNs of individuals with ASD relative to TD individuals. We found that average edge curvature can effectively distinguish the whole-brain functional connectivity of individuals in the ASD and TD groups. Additionally, we studied the differences in node curvature between the two groups and identified specific regions in the brain with atypical functional connectivity in ASD. Notably, we found that brain regions with altered FRC in functional connectivity networks of individuals with ASD, were also associated in the fMRI literature to those cognitive domains known to be affected in ASD. Further, we observed an overlap between the set of regions with altered FRC in functional connectivity networks of individuals with ASD, and those brain regions whose non-invasive stimulation in TMS/tDCS experiments resulted in improvement of ASD-related symptoms.

We acquired rs-fMRI scans of 1112 participants as provided by the ABIDE-I project (Di Martino et al., 2014). The large sample size of the ABIDE-I dataset offers substantial statistical power, thereby increasing the reliability of the reported results (Di Martino et al., 2014; Hull et al., 2017; Lord et al., 2020). We preprocessed each scan using the CONN functional connectivity toolbox (Whitfield-Gabrieli and Nieto-Castanon, 2012), implementing thorough quality assessment (QA) checks both before and after preprocessing. For each participant, we generated a 200 × 200 functional connectivity (FC) matrix using Schaefer atlas (Schaefer et al., 2018) and constructed FCNs with a broad range of edge densities using a maximum spanning tree (MST) followed by sparsity-based thresholding. MST-based network construction is particularly useful for network analyses since it ensures the resulting network is always connected. Similar network construction approaches involving spanning trees have previously been used for financial networks (Samal et al., 2021; Sandhu et al., 2016) and brain FCNs (Achard et al., 2012).

After comparing the average edge curvatures of the FCNs in the ASD and TD groups, we found reduced average FRC and average ORC in individuals with ASD. Similar analysis using standard network measures revealed reduced average clustering coefficient, reduced modularity, reduced average path length, reduced average node betweenness centrality, increased global efficiency and reduced average local efficiency. All the standard network measures except node betweenness centrality have previously been used to study brain-wide changes in functional connectivity in ASD (Harlalka et al., 2018; Itahashi et al., 2014; Rudie et al., 2013), and our results are in agreement with previous findings. However, the changes in graph Ricci curvatures have not previously been studied for FCNs in ASD. Our results illustrate the sensitivity of graph Ricci curvatures, especially FRC, in discriminating the resting state FCNs of individuals with ASD compared to TD.

After comparing the node curvatures of the FCNs in the ASD and TD groups, we identified 83 brain regions that are significantly different in FRC and 14 brain regions that are significantly different in ORC between the two groups. FRC and ORC identify 5 common regions. Moreover, we found that these regions are bilaterally symmetrical and mainly concentrated in 3 RSNs namely, default network, somatomotor network and salient ventral attention network. Previously, Farooq *et al*. (Farooq et al., 2019) have used ORC to compare structural connectivity networks of individuals with ASD relative to TD, and showed that regions with significant difference in ORC are present in visual, dorsal attention, ventral attention areas and temporal lobe. Our results from comparing ORC of resting state FCNs in ASD reveal regions in visual network, dorsal attention network, salient ventral attention network, and additional regions in default network, somatomotor network and limbic network.

We undertook two analyses to assess the agreement of our results with relevant neuroimaging literature on ASD. First, we performed meta-analysis decoding, based on the fMRI literature, to determine if those brain regions with altered FRC in individuals with ASD, were also associated to those cognitive domains that are known to be affected in ASD. We found that brain regions with altered FRC were associated to the cognitive domains of social cognition, memory, movement and language. Farooq *et al*. (Farooq et al., 2019) used ORC to compare structural connectivity networks of individuals with ASD relative to TD, and showed that regions with significant difference in ORC are related to semantic memory, socially relevant memories, emotions and visual perception. Since these results were obtained with structural connectivity networks, it is difficult to compare our results to these. However, each of the cognitive domains suggested by the meta-analysis decoding are known to be affected in ASD (Bhat, 2021; Davidson and Ellis Weismer, 2017; Ellis Weismer et al., 2010; Grace et al., 2018; Griffin et al., 2021; Habib et al., 2019; Kasari and Patterson, 2012; Kristen et al., 2014; Ming et al., 2007; Pickles et al., 2009; Senju, 2012; Solomon et al., 2016; Zampella et al., 2021). Hence, these results suggest that FRC captures atypical connectivity of clinically relevant brain regions underlying ASD. To our knowledge, this is also the first instance of using a meta-analysis decoding approach as a form of validation of results from a graph-theoretic analysis of brain functional connectivity networks.

In addition to meta-analysis decoding based on the fMRI literature, we assessed the agreement of our results with those of TMS/tDCS experiments involving individuals with ASD. We found that brain regions with altered FRC in individuals with ASD overlap with those brain regions whose non-invasive stimulation with TMS/tDCS have been reported to result in improvement of ASD-related symptoms. We further note that the set of regions with altered values of other node-based network measures, *e.g.* clustering coefficient, which overlap with those regions identified by non-invasive stimulation, are a subset of the set of regions identified by FRC. To our knowledge, this is the first instance of using results from TMS/tDCS experiments as a form of validation of results from a graph-theoretic analysis of brain functional connectivity networks. Just as with the comparison of our node-level FRC results against the fMRI literature, the comparison to results from these TMS/tDCS experiments also suggest that FRC captures atypical connectivity of clinically relevant brain regions underlying ASD. Further, FRC might capture atypical connectivity not captured by other node-level network measures such as clustering coefficient. These results commend the use of graph Ricci curvatures as a source of hypotheses about clinically relevant brain regions underlying ASD, which can then be tested by stimulating these regions with non-invasive technologies, *e.g.* TMS (Downar et al., 2016; Lynch et al., 2018; Sale et al., 2015).

To sum up, we find that geometric notions of graph Ricci curvature can be effectively used to determine global and node-level changes in functional connectivity networks of individuals with ASD. Importantly, we present two forms of validation, respectively based on the fMRI and TMS/tDCS literature, to suggest that graph Ricci curvatures, particularly FRC, are sensitive to atypical functional connectivity of clinically relevant brain regions underlying ASD. The methods used in the present work could further be applied to study functional connectivity networks in other atypical populations. Additionally, since graph Ricci curvatures are fundamentally defined on edges, future studies could be aimed at devising edge-based methods to analyze brain functional or structural connectivity.

### Limitations of this study

There was a significant difference in the IQ scores between the ASD and TD groups, which introduces a potential confound to our analysis. Previous studies involving graph-theoretic analysis of rs-fMRI scans in the ABIDE dataset have matched the groups on IQ scores (Harlalka et al., 2018; Keown et al., 2017; Lee et al., 2017). However, we choose to include all subjects in our analysis rather than sub-selecting ASD subjects according to IQ, since sub-selecting would make ASD cohort less representative and the results of our analyses would be less generalizable to the typical ASD population (Dennis et al., 2009). Confining the analysis to IQ-matched ASD subjects would include only high-functioning ASD subjects in the analyses, hence, our results would not be generalizable to ASD subjects whose cognitive functioning is more severely affected. In addition, we have not included IQ as a covariate while comparing the global and local network measures across groups. However, Dennis *et al*. (Dennis et al., 2009) have shown that using IQ as a matching variable or covariate during studies of neurodevelopmental disorders could lead to anomalous findings about neurocognitive function.

## STAR METHODS

### Resource Availability

#### Lead contact

Further information and requests for resources should be directed to and will be fulfilled by the lead contact.

#### Materials availability

This study did not generate new unique materials.

### Method details

In this section, we describe the methodology used to construct the resting-state functional connectivity networks (FCNs) of individuals with autism spectrum disorder (ASD) and typically developing (TD) individuals, from raw resting-state functional MRI (rs-fMRI) images acquired from the Autism Brain Imaging Data Exchange I (ABIDE-I) project (Di Martino *et al*., 2014). Note that TD individuals are the healthy subjects. First, raw rs-fMRI data were spatially and temporally preprocessed using the CONN functional connectivity toolbox (Whitfield-Gabrieli and Nieto-Castanon, 2012). Second, we parcellated the brain into 200 distinct regions of interest (ROIs) or nodes using the Schaefer atlas (Schaefer *et al*., 2018) and a 200 × 200 functional connectivity (FC) matrix was generated for each subject. Third, we filtered the FC matrix using a maximum spanning tree (MST) based approach followed by sparsity-based thresholding to construct FCNs for each subject.

#### Participants and imaging dataset

From the ABIDE-I project (Di Martino *et al*., 2014), we obtained raw rs-fMRI and anatomical data for 1112 participants (age range = 7-64 years, median = 14.7 years), comprising 539 individuals with ASD and 573 age-matched TD individuals. ABIDE-I project is an international effort by 17 imaging sites that have collectively shared rs-fMRI, anatomical and phenotypic data. Further details such as MRI modalities and scan parameters are available on the ABIDE website.

#### Quality assessment and exclusion criteria before preprocessing

We used the following criteria to exclude subjects in ABIDE-I from this study. First, the subjects with missing anatomical or functional files were excluded. Second, all subjects from the imaging site Stanford were excluded as it is the only site with spiral image acquisition protocol. Third, all subjects from the imaging site Leuven-1 were excluded due to unknown repetition times for the functional scans. Fourth, to assess the quality of the raw images in ABIDE-I, we have used the information on raters’ decisions available from the Preprocessed Connectome Project (PCP) (Cameron *et al*., 2013), and the subjects whose raw image quality was described as ‘fail’ by both the raters were excluded. Note that we did not exclude the subjects based on IQ or match the cohorts for IQ in order to ensure that the results of our analyses are generalizable to the typical ASD population (Dennis *et al*., 2009; Di Martino *et al*., 2014). After removing subjects based on the quality assessment (QA) checks and exclusion criteria described above, we were left with 494 subjects in the ASD group and 520 subjects in the TD group (**Supplementary Table S1**).

#### Raw fMRI data preprocessing

We used the CONN functional connectivity toolbox (Whitfield-Gabrieli and Nieto-Castanon, 2012) to process the rs-fMRI data from ABIDE-I. **Figure 1** is a schematic summarizing the processing pipeline for rs-fMRI data used in this study. We have created a protocol video providing a visual guide to rs-fMRI preprocessing using CONN toolbox which is available at: https://youtu.be/ch7-dOA-Vlo.

##### Spatial preprocessing

We performed motion correction, slice-timing correction, outlier detection, and structural and functional segmentation and normalization. First, the functional images were co-registered to the first scan of the first session. The SPM12 realign and unwarp procedure (Andersson *et al*., 2001) was used to realign and motion correct the images using six rigid body transformation parameters: three translations in x, y and z directions, and three rotations namely pitch, yaw and roll. Second, the SPM12 slice-timing correction procedure (Sladky *et al*., 2011) was used to temporally align the functional images. Third, Artifact Detection Tools (ART)-based outlier detection was performed where acquisitions with framewise displacement greater than 0.5 mm or global BOLD signal changes greater than 3 standard deviations were marked as outliers. Fourth, segmentation and normalization (Ashburner and Friston, 2005) was carried out to normalize the images into the standard Montreal Neurological Institute (MNI) space, and then, segment the brain into grey matter, white matter and cerebrospinal fluid (CSF) areas. Raw T1-weighted volume of the anatomical image and mean BOLD signal of the functional images were used as reference in this step. Subjects with bad image quality and signal dropouts in their scans or subjects with registration or normalization errors were excluded from further analysis.

##### Denoising

After the spatial preprocessing of the raw rs-fMRI scans, the BOLD time-series associated with each voxel was extracted using the CONN toolbox. Next, we performed temporal preprocessing or denoising using the CONN toolbox to further reduce physiological or motion effects from the BOLD time-series. First, we implemented anatomical component-based noise correction procedure (aCompCor), to simultaneously remove 5 potential noise components (Chai *et al*., 2012) each from white matter and CSF areas, 12 potential noise components from estimated subject motion parameters and their associated first-order derivatives (Friston *et al*., 1996), and 1 noise component from each of the identified outlier scans (scrubbing) (Power *et al*., 2014) in a single linear regression step. Second, a high-pass filtering was performed to remove temporal frequencies below 0.008 Hz from the BOLD time-series.

#### Quality assessment and exclusion criteria after preprocessing

After preprocessing the raw fMRI data, we applied the following criteria to exclude participants from the analysis. Subjects were excluded if the FC distribution deviated significantly from normal distribution, or if the FC distribution showed noticeable distance dependence (Ciric *et al*., 2017). We additionally excluded subjects that showed a noticeable correlation between quality control (QC) variables and FC values, or if the QC-FC correlations showed a noticeable distance dependence (Ciric *et al*., 2017). After removing subjects based on these exclusion criteria, we were left with 395 subjects in the ASD group and 425 subjects in the TD group (**Supplementary Table S1**). The FC matrices of these remaining 820 subjects were used for network analysis. The demographic and clinical information for these subjects from ABIDE-I included in our study is summarized in **Table 1**.

#### Atlas-based definition of nodes and functional connectivity

A widely-used approach for defining nodes in functional connectivity networks (FCNs) is to group closely related neighboring voxels into cortical parcels, in order to obtain nodes with interpretable neurobiological meaning (Luppi and Stamatakis, 2021). Furthermore, the use of brain parcellations also reduces the computational load of further analyses. In this study, we used a predefined cortical parcellation atlas by Schaefer *et al*., (2018), which is based on a gradient-weighted Markov random field approach. While the Schaefer atlas is available at multiple resolutions, we considered the resolution that parcellates the brain into 200 distinct regions of interests (ROIs) wherein each hemisphere comprises 100 ROIs. In this parcellation, each ROI belongs to one of seven resting state networks (RSNs), namely, ‘visual’, ‘somatomotor’, ‘dorsal attention’, ‘salient ventral attention’, ‘limbic’, ‘control’, and ‘default’. Using the CONN toolbox, the time series of each ROI was computed as the average of the time series of all the voxels that it contains. Subsequently, Pearson correlation coefficient between the time series of every pair of ROIs was calculated in the CONN toolbox, which resulted in a 200 × 200 FC matrix for each subject.

#### Construction of Sparsity-based functional connectivity networks

In the preceding subsection, we described the FC matrix which is a correlation matrix that can be represented as a complete, weighted and undirected graph wherein the ROIs correspond to the nodes and the weights of edges are given by the correlation values between ROIs. The construction of the FCN from the FC matrix of a subject includes two steps, namely maximum spanning tree (MST) construction and sparsity-based thresholding. First, to extract the most important edges from the FC matrix, we constructed its MST using Kruskal’s algorithm (Kruskal, 1956). The MST is a spanning tree of the weighted graph with maximum edge weight. Note that the MST for a weighted graph with *n* nodes is an acyclic graph (more precisely, a tree) with (*n*-1) edges which is always connected. Second, we used sparsity-based thresholding, wherein edges are iteratively added to the MST in decreasing order of their correlation values, until a resulting network with the desired sparsity was obtained. Further, the resulting network with desired sparsity was binarized by ignoring the edge weights before proceeding to compute the network properties (Achard and Bullmore, 2007; Rudie *et al*., 2013).

Evidently, this choice of MST construction followed by sparsity-based thresholding to generate the FCNs ensures that the constructed networks for different subjects are connected and have the same number of edges. Such networks enable direct mathematical comparison of global and local network properties across subjects (Bassett *et al*., 2012; Rudie *et al*., 2013; Xu *et al*., 2016). We remark that this choice of MST followed by sparsity-based thresholding to construct FCNs from rs-fMRI images has been used earlier by Achard *et al*., (2012).

As there is no rationale for using a specific graph density, previous studies (Rudie *et al*., 2013; Itahashi *et al*., 2014; Harlalka *et al*., 2018) have explored network properties across a range of graph densities. In this work, we have studied the network properties over a wide range of graph densities between 0.02 or 2% edges and 0.5 or 50% edges, with an increment of 0.01 or 1% edges. Thus, for each of the 820 subjects from ABIDE-I considered in this study, we have constructed 49 unweighted and undirected networks. In other words, we have generated 820 × 49 FCNs for 820 subjects across 49 graph densities or thresholds for this study, and the constructed networks are made publically available via a GitHub repository.

### Network based analysis and post-hoc analyses

In this section we describe the methodology used to analyze and compare the resting-state FCNs of individuals with ASD and TD individuals constructed as mentioned in the preceding section. First, we performed global and node-level network analysis to compare the FCNs in the ASD group and the TD group. Second, we used Neurosynth-based meta-analysis decoding (Yarkoni *et al*., 2011; Williams *et al*., 2021) to assess the agreement of the results of our node-level network analysis against results reported in a large database of fMRI studies, and studied the relationship between node-level network measures and relevant scores of symptom severity in ASD. Third, we assessed the agreement of the results of our node-level network analysis against results reported in non-invasive brain stimulation (NIBS) studies with Transcranial Magnetic Stimulation (TMS) and transcranial Direct Current Stimulation (tDCS).

#### Global and node-level network analysis

As mentioned in preceding subsection, we constructed 49 unweighted and undirected networks with varying sparsity from the FC matrix corresponding to each subject, and thereafter, each of the 49 networks for a subject was characterized by computing discrete Ricci curvatures and other network properties. Specifically, we have focused here on two discrete Ricci curvatures, namely Forman-Ricci curvature (FRC) (Forman, 2003; Sreejith *et al*., 2016; Samal *et al*., 2018) and Ollivier-Ricci curvature (ORC) (Ollivier, 2007). Notably, the two discrete Ricci curvatures are naturally defined for edges in a network and capture different aspects of the classical Ricci curvature (Samal *et al*., 2018). Moreover, we have also explored here several standard global network measures including average clustering coefficient, modularity (Blondel *et al*., 2008), average shortest path length, average node betweenness centrality, global efficiency (Latora and Marchiori, 2001) and average local efficiency (Latora and Marchiori, 2001). In **Supplementary Information**, we describe the different global and local network measures employed here to characterize the FCNs.

To compare the global properties of the FCNs across the two groups (ASD versus TD), we first computed the average FRC of edges, average ORC of edges and seven other global network measures (including average clustering coefficient, modularity, average shortest path length, average node betweenness centrality, global efficiency and average local efficiency), for each of the 820 × 49 networks corresponding to the FC matrices of 820 subjects across 49 graph densities. To compare the node-level properties of the FCNs across the two groups (ASD versus TD), we computed the node FRC and node ORC for each of the 200 nodes in each of the 820 × 49 networks corresponding to the FC matrices of 820 subjects across 49 graph densities. Note that the node Ricci curvature is defined as the sum of edge Ricci curvatures for the edges incident on that node (Samal *et al*., 2018) (see **Supplementary Information**). Additionally, we computed two standard network measures, namely node clustering coefficient and node betweenness centrality.

The computer codes for FRC and ORC are made publically accessible via a GitHub repository. The other global network measures mentioned above for FCNs were computed using the Python package NetworkX (Hagberg, Schult and Swart, 2008). Furthermore, the statistical tests were performed in Python packages SciPy (Yirtanen et al., 2020) and statsmodels (Seabold and Perktold, 2010).

#### Neurosynth meta-analysis decoding

We used Neurosynth meta-analysis decoding (Yarkoni *et al*., 2011, Williams *et al*., 2021) to determine if the brain regions showing significant between-group differences in node-level network measures have been found in previous fMRI studies to be associated to cognitive domains impaired in ASD, *e.g.* social cognition. Corresponding to each node-level network measure studied here, we identified a set of nodes (ROIs) that showed significant differences between the ASD and TD groups. For a set of nodes with significant between-group differences for a network measure, we used Neurosynth meta-analysis tool to find terms related to cognition, perception and behavior corresponding to the centroid coordinates of each ROI in the set. Further, we partitioned the set of identified ROIs which show significant between-group differences, by the 7 RSNs in the Schaefer atlas, and thereafter, the frequency counts of the terms associated with the subset of identified ROIs in a particular RSN were calculated and the statistical significance of these frequency counts was determined. This was done for each RSN separately, to identify those terms selectively associated with each of the 7 RSNs.

After identifying those cognitive domains associated to the brain regions with significant between-group differences in node-based network measures, we performed a post-hoc correlation analysis to measure the strength of the linear relationship between the values of the node-based network measure for each of the brain regions, and clinical scores related to symptom severity of the identified cognitive domains. We performed this analysis just for the ASD group. Specifically, we chose two clinical scores based on the Autism Diagnostic Interview-Revised (ADI-R) scoring (Lord, Rutter and Le Couteur, 1994), which are: (i) ADI-R verbal and (ii) ADI-R social.

#### Literature search for non-invasive brain stimulation studies in ASD

We performed a literature search to identify brain regions whose non-invasive stimulation were reported to result in improvement of ASD-related symptoms. We first performed the literature search, to identify scientific papers reporting the effect of non-invasive brain stimulation (NIBS) on core symptoms of ASD, and then used results reported in these papers to identify those brain regions whose stimulation resulted in positive behavioral and cognitive outcomes. **Figure 5** summarizes the workflow we employed to collect and classify the eligible articles. We used PubMed to perform the literature search. The search query to PubMed reflected diagnosis of interest including ‘autism spectrum disorder’, ‘Asperger’s syndrome’, ‘autism’ and three major brain stimulation methodologies including ‘Transcranial magnetic stimulation’, ‘TMS’, ‘transcranial direct current stimulation’, ‘tDCS’, ‘transcranial alternating current stimulation’, ‘tACS’. The search was performed in October 2021 and the exact details regarding the search query are provided in **Table 2**. The PubMed search returned 235 articles.

We followed a three-stage procedure to further refine the list of 235 articles returned by the PubMed search. First, we checked for papers missing from the corpus generated by the PubMed search, by scanning review articles on the use of NIBS methods to study ASD. We also searched these review articles for potential databases of NIBS experiments on ASD. Second, we filtered the articles based on title and abstract, based on relevance. We defined relevance according to the following criteria. The inclusion criteria were: (1) studies on ASD populations, (2) studies that have used NIBS techniques namely, TMS (and its variants such as rTMS), tDCS and tACS, (3) studies that have investigated the effect of NIBS on the core behavioral and cognitive symptoms of ASD, and (4) studies that are peer-reviewed. The exclusion criteria were: (1) review articles, (2) articles presented in languages other than English, (3) studies that did not perform NIBS, (4) studies that investigate new protocols for NIBS, (5) studies that report no positive effects in ASD symptoms post NIBS, (6) studies whose target areas for NIBS were not clearly reported, and (7) articles without access to full-text. Third, we classified the articles based on their stimulation technique (TMS/ tDCS/ tACS) and checked the full text of the articles for relevance, according to the same criteria as above. This process yielded 19 eligible articles for TMS, 12 eligible articles for tDCS and zero articles for tACS.

We identified Barahona-Corrêa *et al*. (Barahona-Corrêa *et al*., 2018) as a database of TMS studies in ASD published before 2018, with data collection guided by preferred reporting items for systematic reviews and meta-analysis (PRISMA) (Moher *et al*., 2009). Similarly, we identified García-González *et al*. (García-González *et al*., 2021) as a database of tDCS studies in ASD published before August 2019, also guided by PRISMA data collection. We utilized the data presented in these two databases along with the data that we extracted from the eligible articles in our corpus, such as author, publication year, DOI, number of participants, gender distribution, mean age, intellectual abilities, stimulation methodology and parameters, target areas, stimulation schedule, behavioral and cognitive outcome measures, behavioral and cognitive results, and any adverse reactions for the experiment group and the control group (if applicable). All the data collected are provided as **Supplementary Tables S5** and **S6**. From these data, we identified the set of brain regions whose stimulation using these NIBS methods on individuals with ASD resulted in positive cognitive and behavioral outcomes.

#### Estimating overlap between regions identified in NIBS studies and node-level network analysis

We estimated the overlap between the sets of regions identified from literature search of NIBS studies and the sets of regions revealing ASD-related differences in node-level network measures. The target areas described in the NIBS studies were cortical regions in the brain that are specified by their respective Brodmann areas (Strotzer, 2009) while we identified node-level differences in areas of the Schaefer 200 atlas. We used the MRIcron tool (Rorden, Karnath and Bonilha, 2007) to map each of the Brodmann areas to Schaefer ROIs, by identifying the Brodmann area encompassing the MNI centroid coordinates of each Schaefer ROI (Lynch *et al*., 2018). The mapping from Schaefer ROIs to the Brodmann areas is presented in **Supplementary Table S7**. Next, we compiled the set of Brodmann areas that serve as target areas from the eligible NIBS experiments and have shown a positive outcome, either behavioral or cognitive, as a result of stimulating that region. We then identified the set of Schaefer ROIs that were mapped to these Brodmann areas. From this set of Schaefer ROIs, we found the subset that yielded significant ASD-related differences according to the graph Ricci curvatures namely, FRC and ORC, as well as for clustering coefficient and node betweenness centrality.

### Quantification and statistical analysis

For the global measures, we evaluated the differences between the two groups across the 49 graph densities in the range 2 - 50% considered in this study by using a two-tailed two-sample t-test. For the node-level measures, we first computed the area under the curve (AUC) for a given node measure across the 49 graph densities considered in this study (Achard and Bullmore, 2007; Itahashi *et al*., 2014). Thereafter, we used a two-tailed two-sample t-test to evaluate the differences between the two groups via AUCs of the node measures for each of the 200 nodes in the network. Further, we measured the relationship between the values of the node-based network measure and the ADI-R scores by computing the partial correlations, with age and gender as covariates. For the Neurosynth meta-analysis decoding, to determine statistical significance of these frequency counts, we calculated the frequency counts of the same terms associated with an equal size set of randomly selected surrogate ROIs, and thereafter, the z-score for the frequency counts of each term associated with the subset of original ROIs was calculated. Subsequently, the z-scores were converted into p-values assuming a normal distribution.

After each of the above-mentioned tests or computations, we used a false discovery rate (FDR) correction (Benjamini and Hochberg, 1995) to correct for multiple comparisons and control the occurrence of false positives. Note that the alpha for these FDR corrections was set to 0.05.

## DATA AND CODE AVAILABILITY

- Functional connectivity matrices and networks generated in our study are deposited on GitHub and are publically available as of the date of publication. URL: https://github.com/asamallab/Curvature-FCN-ASD
- All original code to compute the curvature measures is deposited on GitHub and is publically available as of the date of publication. URL: https://github.com/asamallab/Curvature-FCN-ASD
- Any additional information required to reproduce this work is available from the lead contact upon request.

## AUTHOR CONTRIBUTIONS

P.E. and Y.Y. contributed equally to this work. A.S., P.E., Y.Y. and N.W. designed research. P.E. and Y.Y. performed the computations. P.E., Y.Y., N.W., E.S., J.J. and A.S. analyzed data and wrote the paper. P.E. and Y.Y. generated the figures. A.S. supervised the project.

## ACKNOWLEDGEMENTS

We would like to thank Linda Geerligs for discussions. A.S. would like to thank Max Planck Society, Germany for research support via a Max Planck Partner Group in Mathematical Biology. E.S. and J.J. acknowledge support from the German-Israeli Foundation (GIF) grant number I-1514-304.6/2019.

## SUPPLEMENTARY INFORMATION

Supplementary information including Supplementary Methods, Supplementary Figures and Supplementary Tables accompanies the paper.

