## Supplementary Information for "Graph Ricci Curvatures Reveal Atypical Functional Connectivity in Autism Spectrum Disorder"

### Supplementary Methods and Supplementary Figures for Graph Ricci Curvatures Reveal Atypical Functional Connectivity in Autism Spectrum Disorder

Pavithra Elumalai,[1, *] Yasharth Yadav,[1, 2, *] Nitin Williams,[3, 4, †] Emil Saucan,[5] Jürgen Jost,[6, 7] and Areejit Samal[1, 8, ‡]

[1]*The Institute of Mathematical Sciences (IMSc), Chennai, India*
[2]*Indian Institute of Science Education and Research (IISER), Pune, India*
[3]*Helsinki Institute of Information Technology, Department of Computer Science, Aalto University, Finland*
[4]*Department of Neuroscience & Biomedical Engineering, Aalto University, Finland*
[5]*Department of Applied Mathematics, ORT Braude College, Karmiel, Israel*
[6]*Max Planck Institute for Mathematics in the Sciences, Leipzig, Germany*
[7]*The Santa Fe Institute, Santa Fe, NM, USA*
[8]*Homi Bhabha National Institute (HBNI), Mumbai, India*

[*] These authors contributed equally to this work

#### SUPPLEMENTARY METHODS

Each resting state functional connectivity network (FCN) constructed in this work can be represented as an unweighted, undirected and connected graph $G = (V, E)$, where $V$ is the set of vertices (or nodes) and $E$ is the set of edges (or links). There are $n = |V|$ number of nodes and $m = |E|$ number of edges in $G$.

##### Graph Ricci curvatures

The notion of Ricci curvature, originally defined for smooth manifolds (Jost 2017), can also be extended to discrete structures such as networks. The classical definition of Ricci curvature is associated to vectors while its network counterpart is associated to edges (Samal *et al.* 2018). Hence, these curvature measures enable an edge-based approach rather than the node-based approach commonly used in network analysis (Eidi *et al.* 2020). Ricci curvature can be defined for discrete objects in such a way that the ensuing discretization retains some of the key properties of the classical Ricci curvature. In this section, we describe two notions of graph Ricci curvature that we have used to study the global and local properties of FCNs in Autism Spectrum Disorder (ASD).

###### *Ollivier-Ricci curvature*

Ollivier (Ollivier 2007, 2009) introduced a discretization of the classical Ricci curvature, which has been extensively used to analyse complex networks (Samal *et al.* 2018, Lin and Yau 2010, Lin *et al.* 2011, Bauer *et al.* 2012, Jost and Liu 2014, Ni *et al.* 2015, Sandhu *et al.* 2015, 2016, Ni *et al.* 2019, Sia *et al.* 2019). Ollivier-Ricci curvature (ORC) captures the volume growth property of classical Ricci curvature.

A simple interpretation of the classical Ricci curvature can be provided as follows. For two points $x$ and $y$ along a tangent vector $\mathbf{v}$ in a Riemannian manifold, consider a ball (or sphere) $B_x$ of radius $\epsilon$ around $x$. If we use parallel transport to move $B_x$ onto a ball $B_y$ of radius $\epsilon$ around $y$, then the average distance between points on $B_x$ and their corresponding points on $B_y$ is given by:

$$\delta\left(1 - \frac{\epsilon^2}{2(n+2)} Ric(\mathbf{v}) + \mathbf{O}(\epsilon^{\mathbf{3}} + \epsilon^{\mathbf{2}}\delta)\right), \tag{S1}$$

where $\delta = d(x, y)$, $\epsilon, \delta \to 0$, and $Ric(\mathbf{v})$ is the Ricci curvature along $\mathbf{v}$. If $Ric(\mathbf{v})$ is negative, then the average distance that the points of the sphere travel is more than the distance from $x$ to $y$. Similarly, if $Ric(\mathbf{v})$ is positive, then the average distance that the points of the sphere travel is less than the distance from $x$ to $y$.

Ollivier's discretization of the classical Ricci curvature uses an arbitrary probability measure around $x$ instead of a ball of radius $\epsilon$ centered at $x$. More precisely, the ORC of an edge $e$ between nodes $i$ and $j$ in $G$ is defined as

$$\mathbf{O}(e) = 1 - \frac{W_1(m_i, m_j)}{d(i, j)}\ , \tag{S2}$$

where $m_i$ and $m_j$ are the discrete probability measures defined on nodes $i$ and $j$, respectively, and $d(i, j)$ is the distance between $i$ and $j$. For an unweighted graph, $d(i, j)$ is defined as the number of edges contained in the shortest path connecting $i$ and $j$. $W_1$ denotes the Wasserstein distance (Vaserstein 1969), which is the trasportation distance between $m_i$ and $m_j$, given by

$$W_1(m_i, m_j) = \inf_{\mu_{i,j} \in \prod(m_i, m_j)} \sum_{(i',j') \in V \times V} d(i', j')\mu_{i,j}(i', j'), \tag{S3}$$

where $\prod(m_i, m_j)$ is the set of probability measures $\mu_{i,j}$ that satisfy

$$\sum_{j' \in V} \mu_{i,j}(i', j') = m_i(i'),\ \sum_{i' \in V} \mu_{i,j}(i', j') = m_j(j'). \tag{S4}$$

The above equation gives all the possibile transportations of measure $m_i$ to $m_j$ and the Wasserstein distance $W_1(m_i, m_j)$ is the minimal cost of transporting $m_i$ to $m_j$. Note that it is important to specify the probability distribution $m_i$ beforehand. In our computations of Ollivier-Ricci curvature (ORC), it is taken to be uniform over the the neighboring nodes of $i$ (Lin *et al.* 2011).

*Forman-Ricci curvature*

Forman's discretization of the classical Ricci curvature (Forman 2003) was originally introduced in the general framework of *weighted CW cell complexes*, and was later defined in the context of complex networks (Samal *et al.* 2018, Sreejith *et al.* 2016, 2017). Forman-Ricci curvature (FRC) captures the geodesic dispersal property of the classical Ricci curvature and it measures the information spread at the ends of edges in a network (Samal *et al.* 2021). A more negative value of FRC for an edge indicates that the information spread across that edge is higher.

Forman's discretization can be extended to undirected networks such that weights can be assigned to both nodes and edges (Sreejith *et al.* 2016). More precisely, for an edge $e$ between nodes $i$ and $j$ in the graph $G$, FRC is defined as

$$\mathbf{F}(e) = w_e \left( \frac{w_i}{w_e} + \frac{w_j}{w_e} - \sum_{e_i \sim e,\ e_j \sim e} \left[ \frac{w_i}{\sqrt{w_e w_{e_j}}} + \frac{w_j}{\sqrt{w_e w_{e_j}}} \right] \right) \tag{S5}$$

where $w_e$ denotes the weight of the edge $e$, $w_i$ and $w_j$ denote the weights associated with the nodes $i$ and $j$, respectively, $e_i \sim e$ and $e_j \sim e$ denote the set of edges incident on nodes $i$ and $j$, respectively, after excluding the edge $e$. For unweighted graphs, all nodes and edges in $G$ are assigned weight equal to 1. Thus, the expression for FRC reduces to

$$\mathbf{F}(e) = 4 - deg(i) - deg(j) \tag{S6}$$

where $deg(i)$ and $deg(j)$ are the degrees of nodes $i$ and $j$, respectively.

Samal *et al.* (Samal *et al.* 2018) have introduced a modified version of FRC for complex networks, known as *augmented Forman-Ricci curvature*, where a graph is constructed as a two-dimensional simplicial complex. A two-dimensional simplicial complex comprises vertices, edges and triangular faces. Thus, augmented FRC accounts for cycles of length 3 in a graph while neglecting cycles of length 4 or higher. For a graph $G$ where weights are assigned to vertices, edges and triangular faces, the augmented FRC of an edge $e$ is defined as

$$\mathbf{F}^{\#}(e) = w_e \left[ \left( \sum_{e<f} \frac{w_e}{w_f} + \sum_{v<e} \frac{w_v}{w_e} \right) - \sum_{\hat{e} \| e} \left| \sum_{\hat{e}, e<f} \frac{\sqrt{w_e \cdot w_{\hat{e}}}}{w_f} - \sum_{v<\hat{e}, e} \frac{w_v}{\sqrt{w_e \cdot w_{\hat{e}}}} \right| \right] \tag{S7}$$

where $w_e$ is the weight of edge $e$, $w_v$ is the weight of vertex $v$, $w_f$ is the weight of triangular face $f$. $\sigma < \tau$ means that the $\sigma$ is a lower dimensional face of $\tau$, $\|$ means that two cells are parallel, i.e. they either share a higher dimensional face or a lower dimensional face, but not both. If the graph $G$ is unweighted, then all the vertices, edges and triangular faces are assigned weight equal to 1, and the augmented FRC reduces to the following simple expression

$$\mathbf{F}^{\#}(e) = \mathbf{F}(e) + 3m \tag{S8}$$

where $\mathbf{F}(e)$ is the FRC of the edge $e$ and $m$ is the number of triangular faces that contain $e$. In this study, we refer to augmented FRC as FRC throughout the main text.

**Standard network measures**

The standard network measures employed in the present work are defined for an unweighted, undirected, and connected graph $G = (V, E)$. The graph $G$ can also be represented by an $n \times n$ adjacency matrix $\mathbf{A}$, with its elements defined as $A_{ij} = 1$ if nodes $i$ and $j$ are adjacent, and $A_{ij} = 0$ otherwise.

- The *clustering coefficient* measures the tendency of nodes in a graph to form triangles with their neighbors. For a node $i$ in the graph $G$, clustering coefficient is defined as:

  $$C_i = \frac{2}{k_i(k_i - 1)} \sum_{j,k} (A_{ij} A_{ik} A_{jk})^{1/3} \tag{S9}$$

  where $j$ and $k$ are the neighbors of node $i$ and the summation is over all neighboring pairs. The *average clustering coefficent* of $G$ is the average of the clustering coefficients of the individual nodes in $G$.

- A common property of many networks is the tendency to exhibit a modular structure, where the set of nodes in a network can be partitioned into subsets of densely connected nodes. *Modularity* measures the density of intra-module edges compared to inter-module edges. For the graph $G$, modularity is defined as (Girvan and Newman 2002, Blondel *et al.* 2008)

$$Q = \frac{1}{2m_w} \sum_{i \neq j \in V} [\, A_{ij} - \frac{s_i s_j}{2m_w}] \delta(c_i, c_j) \tag{S10}$$

where $s_i$ and $s_j$ give the sum of weights of edges attached to nodes $i$ and $j$, respectively, $c_i$ and $c_j$ are the communities of $i$ and $j$, respectively, and $m_w$ is the sum of all edge weights. Since $G$ is an unweighted graph, all edges in $G$ are assigned weight equal to 1.

- The *shortest path length* $d(i,j)$ between any two nodes $i$ and $j$ in $G$ is defined as the number of edges contained in the shortest path connecting them. *Average shortest path length* is the average of the shortest path length between all pairs of nodes, i.e.,

$$\langle L \rangle = \frac{1}{n(n-1)} \sum_{i,j \in V} d(i,j) \tag{S11}$$

- The *betweenness centrality* of a node measures the extent to which it lies on the shortest path between other nodes. For a node $i$ in $G$, node betweenness centrality is defined as (Freeman 1977)

$$C_b(i) = \sum_{j,k \in V} \frac{\sigma(j,k|i)}{\sigma(j,k)}, \tag{S12}$$

where $\sigma(j,k)$ is the number of shortest paths between $j$ and $k$, and $\sigma(j,k|i)$ is the number of shortest paths between $j$ and $k$ that pass through $i$.

- *Global efficiency* was introduced in order to account for information flow in a network (Latora and Marchiori 2001). It measures the global ability of the network to exchange information. Global efficiency of $G$ is defined as

$$E(G) = \frac{1}{n(n-1)} \sum_{i \neq j \in V} \frac{1}{d_{ij}}. \tag{S13}$$

- Define $G_i$ as the subgraph of the neighbors of node $i$ in $G$. Then the *local efficiency* (Latora and Marchiori 2001) of node $i$, $E(G_i)$ is the efficiency of the subgraph $G_i$. *Average local efficiency* is the average efficiency of local subgraphs in $G$.

$$E_{loc} = \frac{1}{n} \sum_{i \in V} E(G_i) \tag{S14}$$

#### SUPPLEMENTARY FIGURES

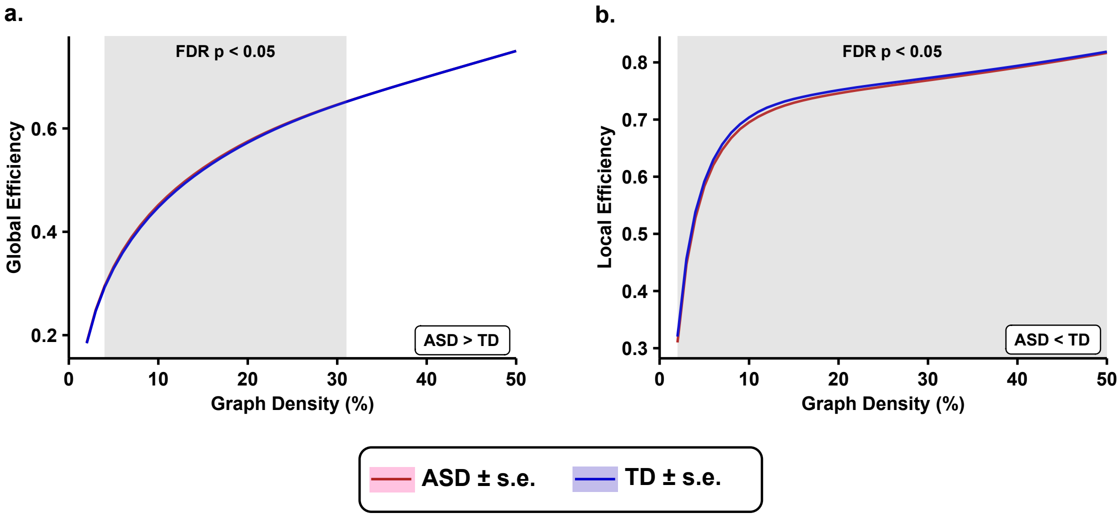


FIG. S1. **Comparison plots related to brain wide changes in functional connectivity networks (FCNs) as captured by network measures between 395 subjects with autism spectrum disorder (ASD) and 425 typically developing individuals (TD)**. Each network measure was compared over a wide range of graph densities between 0.02 (i.e., 2% edges) and 0.5 (i.e., 50% edges), with an increment of 0.01 (i.e., 1% edges). The shaded regions in each plot indicate statistically significant differences ($p < 0.05$, FDR-corrected) between the two groups at the corresponding graph densities on the x-axis. Even though the differences are not explicit from the plots, the directionalities are programmatically verified. (a) Global efficiency is significantly increased in the ASD group across graph densities 4% - 31%. (b) Average local efficiency is significantly reduced in the ASD group across graph densities 2% - 50%.

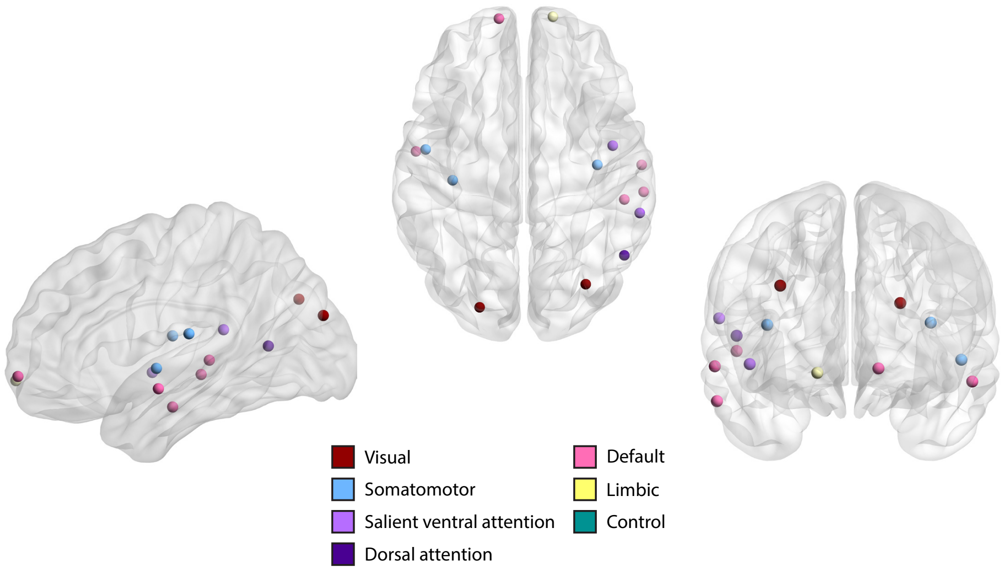


FIG. S2. **Visual representation of 14 nodes or regions in the brain that are significantly different ($p < 0.05$, FDR-corrected) between individuals with autism spectrum disorder (ASD) and typically developing individuals (TD), as captured by Ollivier-Ricci curvature (ORC) of the nodes, related to region specific changes in functional connectivity networks**. The nodes are defined using the Schaefer atlas and each node belongs to one of 7 resting state networks (RSNs) as listed in the figure legend. We find that identified nodes are mainly concentrated within the default network and somatomotor network. This figure was created using BrainNet Viewer (Xia *et al.* 2013).

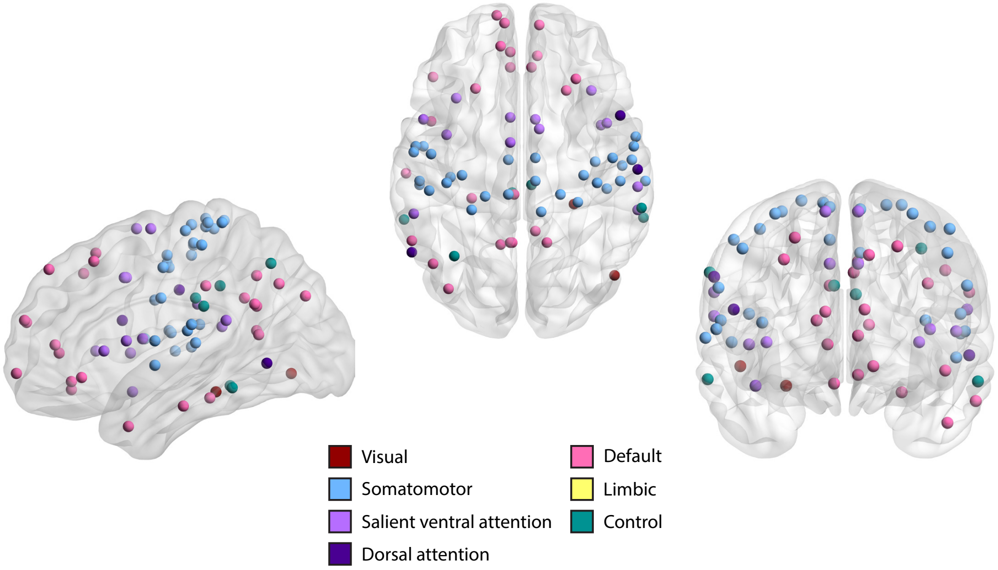


FIG. S3. **Visual representation of 78 nodes or regions in the brain that are significantly different ($p < 0.05$, FDR-corrected) between individuals with autism spectrum disorder (ASD) and typically developing individuals (TD), as captured by clustering coefficient, related to region-specific changes in functional connectivity networks**. The nodes are defined using the Schaefer atlas and each node belongs to one of 7 resting state networks (RSNs) as listed in the figure legend. We find that identified nodes are mainly concentrated within the default network, somatomotor network and salient ventral attention network. This figure was created using BrainNet Viewer (Xia *et al.* 2013).

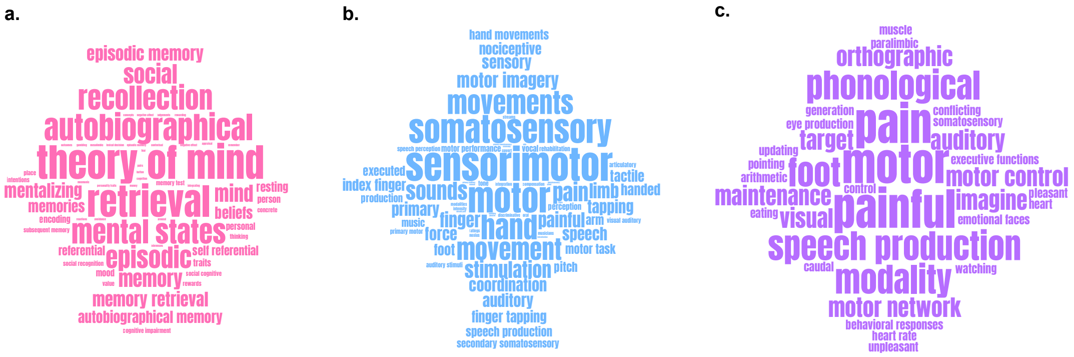


FIG. S4. **The word clouds depicting the behavioral relevance of nodes or regions in different resting state networks (RSNs) that are significantly different ($p < 0.05$, FDR-corrected) between individuals with autism spectrum disorder (ASD) and typically developing individuals (TD) as captured by clustering coefficient of the nodes, relating to agreement of results from node-based network analysis to fMRI literature**. The size of the terms in each word cloud indicates their frequency count. Note that that size of the terms in each word cloud are scaled separately and thus the frequency counts cannot be compared across word clouds. (a) The word cloud corresponding to the nodes in the default network that show significant differences in clustering coefficient. (b) The word cloud corresponding to the nodes in the somatomotor network that show significant differences in clustering coefficient. (c) The word cloud corresponding to the nodes in the salient ventral attention network that show significant differences in clustering coefficient. The word clouds are generated using wordclouds.com.

J. Jost, *Riemannian geometry and geometric analysis*, 7th ed. (Springer Berlin Heidelberg, New York, NY, 2017).
A. Samal, R. P. Sreejith, J. Gu, S. Liu, E. Saucan, *et al.*, Scientific Reports **8**, 8650 (2018).
M. Eidi, A. Farzam, W. Leal, A. Samal, and J. Jost, Theory in Biosciences **139**, 337 (2020).
Y. Ollivier, Comptes Rendus Mathematique **345**, 643 (2007).
Y. Ollivier, Journal of Functional Analysis **256**, 810 (2009).
Y. Lin and S.-T. Yau, Mathematical Research Letters **17**, 343 (2010).
Y. Lin, L. Lu, and S.-T. Yau, Tohoku Mathematical Journal **63**, 605 (2011).
F. Bauer, J. Jost, and S. Liu, Math. Res. Lett. **19**, 1185 (2012).
J. Jost and S. Liu, Discrete & Computational Geometry **51**, 300 (2014).
C. Ni, Y. Lin, J. Gao, X. D. Gu, and E. Saucan, in *2015 IEEE Conference on Computer Communications (INFOCOM)* (IEEE, 2015) pp. 2758–2766.
R. Sandhu, T. Georgiou, E. Reznik, L. Zhu, I. Kolesov, *et al.*, Scientific Reports **5**, 12323 (2015).
R. S. Sandhu, T. T. Georgiou, and A. R. Tannenbaum, Science Advances **2**, e1501495 (2016).
C.-C. Ni, Y.-Y. Lin, F. Luo, and J. Gao, Scientific Reports **9**, 9984 (2019).
J. Sia, E. Jonckheere, and P. Bogdan, Scientific Reports **9**, 9800 (2019).
L. N. Vaserstein, Probl. Peredachi Inf. **5**, 64 (1969).
R. Forman, Discrete and Computational Geometry **29**, 323 (2003).
R. P. Sreejith, K. Mohanraj, J. Jost, E. Saucan, and A. Samal, Journal of Statistical Mechanics: Theory and Experiment **2016**, 063206 (2016).
R. Sreejith, J. Jost, E. Saucan, and A. Samal, Chaos, Solitons & Fractals **101**, 50 (2017).
A. Samal, H. K. Pharasi, S. J. Ramaia, H. Kannan, E. Saucan, *et al.*, Royal Society Open Science **8**, rsos.201734, 201734 (2021).
M. Girvan and M. E. J. Newman, Proceedings of the National Academy of Sciences **99**, 7821 (2002).
V. D. Blondel, J.-L. Guillaume, R. Lambiotte, and E. Lefebvre, Journal of Statistical Mechanics: Theory and Experiment **2008**, P10008 (2008).
L. C. Freeman, Sociometry **40**, 35 (1977).
V. Latora and M. Marchiori, Physical Review Letters **87**, 198701 (2001).
M. Xia, J. Wang, and Y. He, PloS one **8**, e68910 (2013).
